# The plant immune receptors NRG1.1 and ADR1 are calcium influx channels

**DOI:** 10.1101/2021.02.25.431980

**Authors:** Pierre Jacob, Nak Hyun Kim, Feihua Wu, Farid El-Kasmi, William G. Walton, Oliver J. Furzer, Adam D. Lietzan, Sruthi Sunil, Korina Kempthorn, Matthew R. Redinbo, Zhen-Ming Pei, Li Wan, Jeffery L. Dangl

## Abstract

Plant nucleotide-binding leucine-rich repeat receptors (NLRs) regulate immunity and cell death. RPW8 domain-containing “helper” NLRs (RNLs) are required by many “sensor” NLRs. Our crystal structure of the RNL N REQUIREMENT GENE 1.1 (NRG1.1) N-terminal signaling domain resembled that of the resting state plant resistosome-forming HOPZ-ACTIVATED RESISTANCE 1 (ZAR1) and the animal MIXED-LINEAGE KINASE-LIKE (MLKL) cation channel. Active NRG1.1 oligomerized, was enriched in plasma membrane puncta and conferred cytoplasmic Ca^2+^ influx in plant and human HeLa cells. NRG1.1-dependent Ca^2+^ influx and cell death were sensitive to Ca^2+^ channel blockers. Ca^2+^ influx and cell death mediated by NRG1.1 and ACTIVATED DISEASE RESISTANCE 1 (ADR1), another RNL, required conserved negatively charged N-terminal residues. Thus, RNLs apparently form influx channels to directly regulate cytoplasmic [Ca^2+^] and consequent cell death.

**One Sentence Summary:** A specific class of plant immune receptors function as calcium-permeable channels upon activation to induce cell death.

---

Nucleotide-binding leucine-rich repeat receptors (NLRs) are major determinants of immunity (*1*). In plants, two major classes of NLRs exist based on their N-terminal domains: Toll/Interleukin-1 receptor (TIR)-NLRs, (hereafter, TNL) and the coiled-coil (CC)-NLRs (hereafter, CNL) (*2, 3*). All tested TNLs require ENHANCED DISEASE SUSCEPTIBILITY 1 (EDS1) as well as a small subfamily of redundant “helper” NLRs also known as RNLs due to their CC-R [RPW8 (Resistance to Powdery Mildew 8)-like CC] domain (*4-6*). There are two sub-families of RNLs, NRG1 (N REQUIREMENT GENE 1) and ADR1 (ACTIVATED DISEASE RESISTANCE 1), in nearly all flowering plants (*7*).

We demonstrated that an auto-active allele of the *Arabidopsis* RNL NRG1.1 (N REQUIREMENT GENE 1.1, D485V; hereafter DV), mutated in the conserved MHD motif and a common proxy for pathogen-mediated NLR activation, (*8*) was sufficient to trigger cell death *in planta* (Fig. S1). This cell death was independent of the native RNLs NRG1 and ADR1 (*3, 9, 10*) and of the EDS1 signaling module (*6, 10*) in the heterologous host, *Nicotiana benthamiana* (Fig. S1)

We obtained X-ray crystal structures of two mutant NRG1.1 CC-R domains (residues 1-124), K94E/K96E/R99E/K100E/R103E/K106E/K110E [7K(R)/E] and K94E/K96E (2K/E) that diffracted to 3.0 Å and 2.5 Å, respectively (Table S1). These putative surface mutations were required to achieve monodispersion of the protein. Structural homology modeling suggested that the CC-R domains share an N-terminal 4-helical bundle (4HB) with cell death pore forming mammalian mixed-linage kinase domain-like (MLKL) proteins and fungal HET-s/HELO domain proteins (*5, 11, 12*). Similar to predictions, the two mutant structures superimposed well with the 4HBs of the resting-state CC domain of ZAR1 (*13*) and the cation channel-forming domain of MLKL (*14*) (Fig. 1, A to D; Table S2).

**Figure 1.**
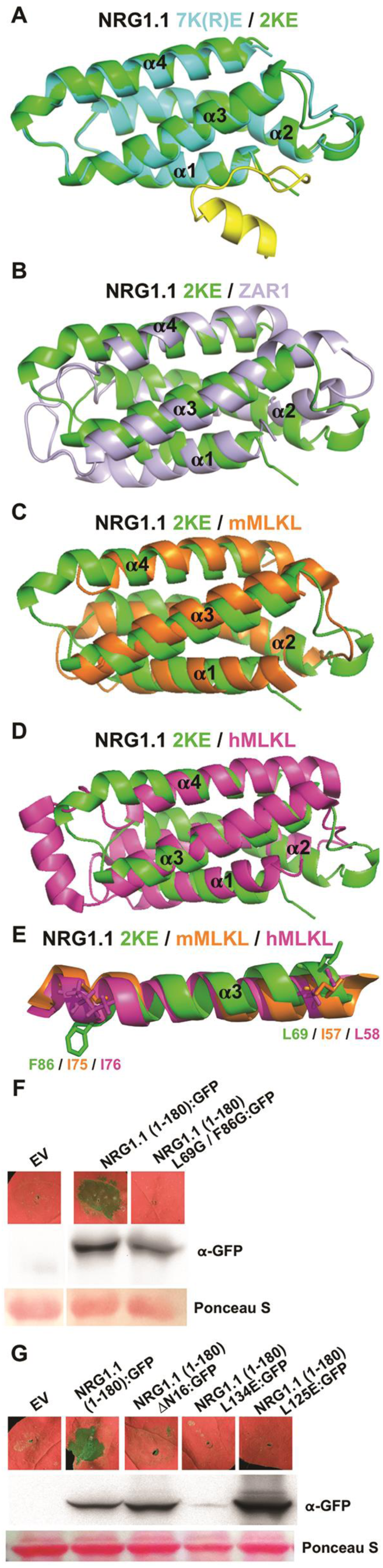
NRG1.1 CC-R resembles ZAR1 and MLKL 4HB. (**A**) Structural overlay of 7K(R)E (PBD: 7L7V) and 2KE (7L7W) with 2KE in green, 7K(R)E 4HB in cyan and 7K(R)E N-terminal region in yellow. (**B**) Structural overlay of 2KE in green and the ZAR1 4HB (6J5V) structure in light purple. (**C**) Structural overlay of 2KE in green and mMLKL 4HB (4BTF) in orange. (**D**) Structural overlay of 2KE in green and hMLKL 4HB (2MSV) in magenta. (**E**) Highlight of the conserved F86/I75/I76 and L69/I57/L58 shown in sticks in the α3 helix of superimposed 2KE, mMLKL and hMLKL 4HB structures, respectively. In planta (*Nicotiana benthamiana; Nb*) phenotypes at 2 days post induction (dpi) and protein accumulation of WT NRG1.1 CC-R and structure-derived mutants (**F** and **G**). EV, empty vector.

The 4HB of MLKL is sufficient to cause cell death and this requires two hydrophobic residues L58 and I76 that maintain the 4HB hydrophobic core (*15*). Structural overlay predicted that the two hydrophobic residues are conserved in NRG1.1 as L69 and F86 (Fig. 1E); mutating them abolished cell death activity of NRG1.1 1-180 (Fig. 1F). In ZAR1, a hydrophobic groove made by α2 and α4B is important for oligomerization and function (*13*). We found two hydrophobic residues L134 and L125 in the homologous region in NRG1.1; mutating them to glutamic acid abolished cell death in NRG1.1 1-180 (Fig. 1G). These data validate the NRG1.1 4HB structure.

Active ZAR1 oligomerizes into a pentamer where the α1 helix of the 4HB re-arranges, flips out and forms a proposed plasma membrane (PM)-penetrating, funnel-like structure (*13, 16*), consistent with models proposed for fungal Het-s/HELO activation (*17*). This first helix was essential for ZAR1 function at the PM (*13*). It was consequently posited that ZAR1 would perturb the PM, but it remains unclear how the postulated funnel-like structure mediates PM permeability, substrate selectivity, cell death and defense activation in plants (*13, 16, 18*). One of the NRG1.1 CC-R structures (7(K)R/E) revealed a potentially flexible N-terminus (residues 1-16; absent in ZAR1; Fig. 1A) that could extend the 4HB α1 helix of a putative funnel in the active NRG1.1 protein. This was disordered in the 2K/E structure. This region was required for NRG1.1 1-180 cell death induction (ΔN16; Fig. 1G).

We introduced several of the structure-derived mutations into the full length NRG1.1 in *cis* with the auto-active DV allele, D485V, because the wildtype resting (WT) NRG1.1 is inactive in the absence of sensor NLR activation (Fig. 2A). We found that the ΔN16, L134E and L125E *cis* mutations each abolished the cell death function of NRG1.1 DV (Fig. 2A). Blue native (BN)- PAGE analyses revealed that active NRG1.1 DV formed high molecular weight complexes whereas the inactive WT, catalytic p-loop (G199A/K200A) and DV p-loop *cis* double mutants did not (Fig 2B). Mutation of L134E, but not of L125E, in *cis*, also abolished NRG1.1 DV oligomer formation (Fig. 2B). Unlike the WT or inactive p-loop mutants, NRG1.1 DV was enriched in the PM fraction while DV L125E showed significantly reduced PM enrichment (Fig. 2C). Although it oligomerized and was enriched at the PM (Fig. S2-4), NRG1.1 DV ΔN16 failed to induce cell death (Fig. 2, A to C).

**Figure 2.**
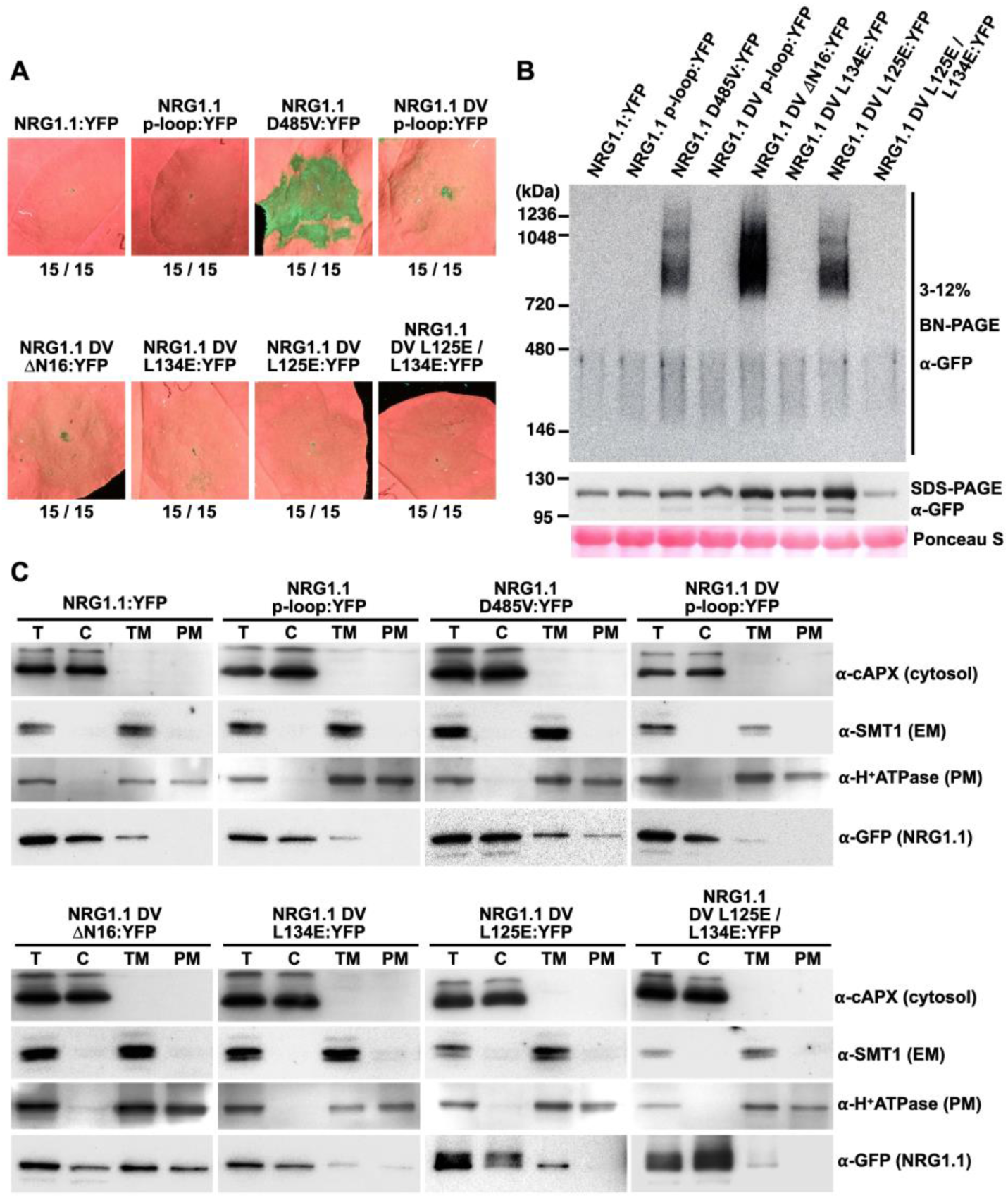
Active NRG1.1 (D485V) oligomerizes on the plasma membrane and triggers cell death. (**A**) In planta phenotypes of NRG1.1 variants in *Nb* at 2 dpi. (**B**) Oligomerization of NRG1.1 DV variants. Accumulation of YFP-tagged NRG1.1 variants were verified by SDS-PAGE- immunoblotting while their oligomeric states were checked by BN-PAGE-immunoblotting in *Nb adr1 nrg1*. (**C**) Activation promotes PM localization. Total proteins (T) extracted from *Nb* expressing NRG1.1:YFP variants were fractionated into cytosolic (C), total membrane (TM) and plasma membrane (PM) fractions and verified by marker proteins: Cytosol, cytosolic ascorbic peroxidase; endoplasmic reticulum membrane (EM), sterol methyltransferase 1; PM, H^+^ATPase.

Confocal microscopy demonstrated that NRG1.1 DV exhibits increased PM localization compared to inactive alleles (Fig. S2-4). Notably, active NRG1.1 DV exhibited puncta on the PM while the NRG1.1 DV p-loop double mutant exhibited many fewer (Fig. S2-4). These PM puncta were also observed for the NRG1.1 DV ΔN16 double mutant but were less common for the missense loss of function alleles, which co-localized more with the ER marker than the PM marker (Fig. S3). Collectively, these results highlight the importance of the NRG1.1 N-terminal 16 residues, which we speculate could extend a putative funnel-like structure by analogy with ZAR1.

These results, particularly the observation that NRG1.1 ΔN16 retains oligomerization and PM puncta formation but loses function, prompted us to investigate the possibility that NRG1.1 forms PM pores and functions as a channel. To observe the sole effects of NRG1.1, we expressed NRG1.1 variants in HeLa cells. If NRG1.1 caused cell death, formed pores, and exhibited channel activity in this evolutionarily distant cellular background, the most parsimonious conclusion would be that it did so autonomously. The alternative hypothesis would require a signaling partner conserved between plants and humans. NRG1.1 DV induced significant cell death at 6 hours post-induction (Fig. S5). This did not occur when WT NRG1.1 was expressed or when NRG1.1 DV activity was suppressed by p-loop, ΔN16, L125E and L134E mutations in *cis* (Fig. S5). Oligomerization and PM localization of the NRG1.1 variants in HeLa cells were similar to the *in planta* results (Fig. S6). Thus, the requirements for NRG1.1 cell death induction are similar in HeLa cells and in plant cells.

We observed the morphology of the dying HeLa cells using scanning electron microscopy. We noted a significantly increased number of PM pores correlated with the cell death activity of NRG1.1 variants (Fig. S7, Fig. 3A-C). NRG1.1 DV expressing cells exhibited 8nm (average) pores in these processed samples (Fig. 3E, F), although this may not represent the actual pore size. This apparent pore size was significantly different from the size of larger pores formed by GASDERMIN-D 1-275 L192D, a partial loss of function mutant of the pore-forming protein GASDERMIN-D that induces cell death and allows high protein accumulation ((*13, 19*), Fig. 3D-F; Fig. S5B). Cell fractionation experiments indicated that NRG1.1 DV was enriched in the PM fraction in HeLa cells whereas the NRG1.1 DV p-loop *cis* double mutant was not (Fig. S6B). Overall, active NRG1.1 DV localized to the PM and its cell death activity was associated with the occurrence of PM pores in HeLa cells.

**Fig3:**
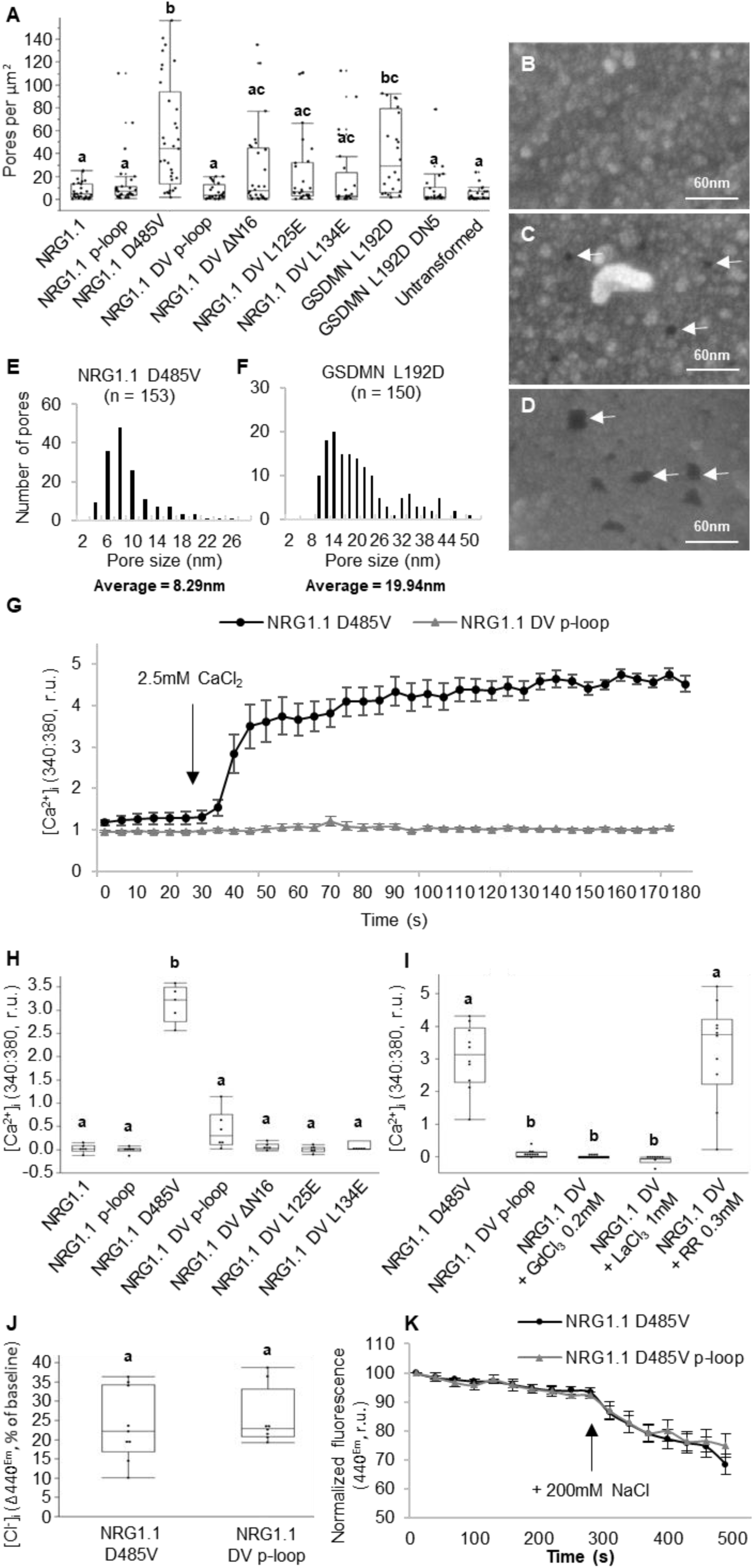
NRG1.1 forms ion channels permeable to Ca^2+^ and not Cl^-^. (**A**) Quantification of apparent PM pores in HeLa cells expressing NRG1.1 variants. (**B**) to (**D**), representative scanning electron micrographs of cells exhibiting apparent PM pores in cell lines expressing NRG1.1 DV p-loop (**B**), NRG1.1 D485V (**C**) or the pore forming protein GSDMN 1-275 L192D (**D**). White arrows indicate the visible PM pores. (**E**) and (**F**) Distribution of the diameter of PM pores visible after NRG1.1 D485V (**E**) or GSDMN L192D (**F**) expression. The average diameters are significantly different (t-test, p-value<0.0001). (**G**) [Ca^2+^]_i_ in NRG1.1 D485V or NRG1.1 D485V p-loop expressing HeLa cells. Black arrow indicates CaCl_2_ addition. (**H**) Ca^2+^ influx in HeLa cells expressing NRG1.1 variants. (**I**) Effect of Ca^2+^ channel blockers LaCl_3_, GdCl_3_ and Ruthenium Red (RR) on NRG1.1 D485V-induced Ca^2+^ influx. (**J**) Intracellular [Cl^-^] accumulation in cells expressing NRG1.1 D485V or NRG1.1 DV p-loop. (**K**) Representative time course experiment showing variation of [Cl^-^]. Black arrow indicates 200mM NaCl addition. Letters indicate statistical significance (ANOVA with Post Hoc Tukey HSD, p- value < 0.01).

Ca^2+^ influx is a hallmark of programmed cell death in both animal and plant kingdoms (*20, 21*), and a requirement for NLR signaling (*20, 22*). We measured Ca^2+^ influx in HeLa cells expressing NRG1.1 DV or the DV p-loop double mutant to see if the increased PM pores affected cytoplasmic Ca^2+^ homeostasis. We observed sustained intracellular Ca^2+^ concentration ([Ca^2+^]_i_) increase specifically in the NRG1.1 DV expressing cells seconds after CaCl_2_ addition (Fig. 3G, Fig. S8). Loss of cell death *cis* mutations ΔN16, L125E or L134E all suppressed NRG1.1 DV-driven Ca^2+^ influx (Fig. 3H, Fig. S8). The general Ca^2+^ influx channel blockers LaCl_3_ and GdCl_3_ but not the Ca^2+^-induced Ca^2+^ release channel blocker ruthenium red (RR) (*23-25*), blocked NRG1.1 DV-driven Ca^2+^ influx (Fig. 3I, Fig. S9). These observations are consistent with NRG1.1 DV forming ion channels that mediate Ca^2+^ influx. We investigated the specificity of NRG1.1 DV-driven ion flux by measuring cytosolic [Cl^-^] using 6-methoxy-N-ethylquinolinium iodide (MEQ)/dihydroMEQ (*26*). There was no difference between the NRG1.1 DV and NRG1.1 DV p-loop expressing cells (Fig. 3, J, K, Fig. S10), indicating that NRG1.1 DV channels do not drive Cl^-^ influx. Thus, we conclude that NRG1.1 forms Ca^2+^- permeable channels in the PM of HeLa cells.

We focused on the first 16 amino acids of NRG1.1 across 334 plant RNL sequences, since NRG1.1 DV ΔN16 retained oligomerization and PM localization but lost Ca^2+^ influx. We observed a pattern of glycine or negatively charged or polar residues separated by 2 to 3 hydrophobic residues (Table S3, Fig. S11). This motif was conserved in the ADR1 clade of RNLs, partially degenerated in the NRG1 clade and further degenerated in CNLs (Fig. -S12). While different from the recently described N-terminal MADA motif (*27*) these two domains share regularly spaced negatively charged residues.

Such residues are critical for ion selectivity and permeability in Ca^2+^ channels (*28, 29*). We therefore targeted the negatively charged residues within the first 16aa of NRG1.1 and ADR1 for structurally conservative but uncharged *cis* mutations (NRG1.1 DV: D3N, E14Q or D3N/E14Q; ADR1: D6N, D11N or D6N/D11N).

We assayed Ca^2+^ influx triggered by these alleles *in planta* using the intracellular Ca^2+^ reporter GCaMP3 in transgenic *N. benthamiana (30*). We confirmed that both NRG1.1 DV-mediated and ADR1 expression triggered [Ca^2+^]_i_ increases *in planta*, accompanied by cell death (Fig. 4A,B). LaCl_3_ and GdCl_3_ abolished cell death (Fig. 4A) and reduced Ca^2+^ influx (Fig. 4B). In both plant (Fig. 4 C-F) and HeLa cells (Fig. 4 G-J) mutations in the RNL-conserved N-terminal motif, especially E14Q (in NRG1.1 DV) and D11N (in ADR1), significantly reduced the rate of Ca^2+^ influx (Fig 4, C, E, G-J; Fig. S13). These mutations also severely reduced cell death induction (Fig. 4, D, F; Fig. S14-S15). These results indicate that the conserved negatively charged residues in the RNL motif are required for appropriate Ca^2+^ influx.

**Figure 4.**
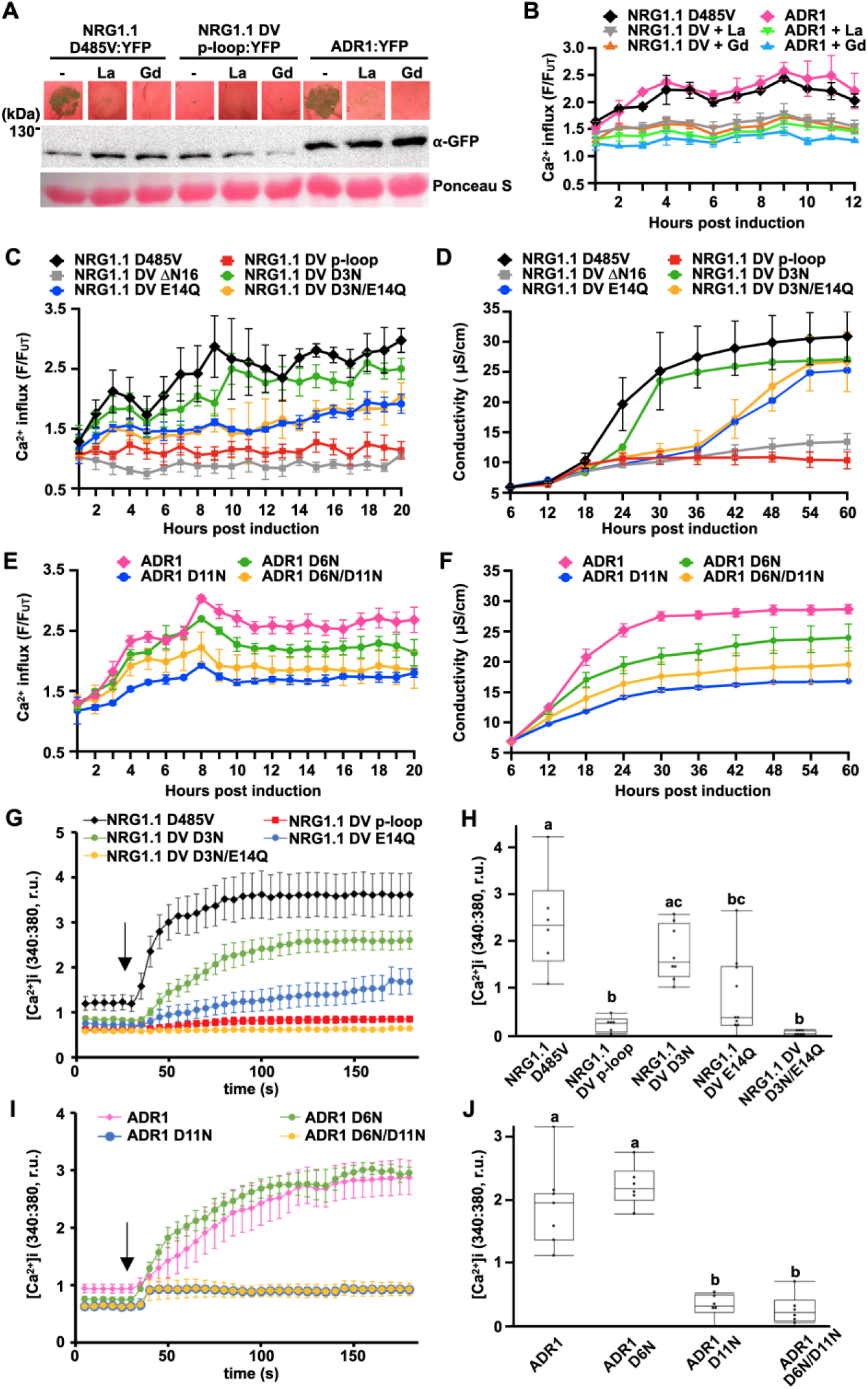
Negatively-charged residues in the conserved RNL motif are required for Ca^2^ influx and cell death. **(A)** In planta phenotypes (2 dpi) of plant RNLs in the presence of Ca^2+^ channel blockers LaCl_3_ (La) and GdCl_3_ (Gd). Accumulation of YFP-tagged RNL variants were verified by SDS-PAGE and-immunoblotting, in *Nb adr1/nrg1*. **(B**), **(C)** and **(E)** Time-course RNL-induced Ca^2+^ influx measurements with GCaMP3 transgenic *Nb*. (**D**) and (**F**) Time-course conductivity measurement depicting RNL triggered cell death in *Nb*. (**G**) to (**J**), [Ca^2+^]_i_ in HeLa cells expressing NRG1.1 DV and variants, (**G**) and (**H**), or ADR1 and variants, (**I**) and (**J)**. Black arrows indicate the addition of CaCl_2_. Letters indicate statistical significance (ANOVA with Post Hoc Tukey HSD, p-value < 0.05). YFP-tagged RNL protein expression in *Nb* and HeLa cells was verified (Fig. S16).

Our data are most consistent with active RNLs functioning as Ca^2+^-permeable channels. The N-terminal 16 residues of NRG1.1 are required for function and Ca^2+^ influx, but not for oligomerization or PM localization. Within this region, a conserved negatively charged residue, E14 in NRG1.1 and D11 in ADR1, is specifically required for Ca^2+^ influx and cell death.

Ca^2+^ signaling is a major regulator of plant immunity (*21, 31*). All TNL immune receptors tested so far require the redundant RNLs of the ADR1 and NRG1 subfamilies (*4*). Thus, we propose that TNL activation induces RNL-dependent Ca^2+^ influx, which triggers defense and/or cell death. CNLs may also directly trigger cytoplasmic Ca^2+^ influx. Among CNLs, at least ZAR1 and RPM1 do not strictly require RNLs for function (*4, 5*). Cell death induced by RPM1 is inhibited by LaCl_3_, GdCl_3_ and RR (*23*). In addition, RPM1 activation correlates with sustained Ca^2+^ influx in plant cells (*22*). The α1 helix in active ZAR1 forms a funnel required for PM interaction that could form PM pores (*13*). ZAR1 E11/E18 form a ring of inward facing E residues in the ZAR1 pentameric funnel and are required for function (*13*), similar to the requirement for negatively charged residues in NRG1.1 and ADR1 that we describe here. A double mutant of E14/E27 in NRG1.1 loses both cell death and defense activation but retains interaction with required co-regulators (*32*). A similar ring of E residues was shown to function in Ca^2+^ binding and Ca^2+^ permeability of voltage-gated Ca^2+^ channels (*28*). It is tempting to speculate that ZAR1 E11, NRG1.1 E14 and ADR1 D11 all form E/D-type rings that act as Ca^2+^ selective filters to achieve fast Ca^2+^ influx and initiate cell death signaling.

## Acknowledgements

We thank Drs. Brian Staskawicz, Raoul Martin (UCB), Julian Schroeder (UCSD) and Dr. Sarah Grant (UNC) for critical reading of the manuscript and the Dangl lab NLR group for helpful discussions of the work. Supported by the National Science Foundation (Grant IOS-1758400 to J.L.D. and IOS-1457257 to Z.-M.P.), the National Institutes of Health (grants GM137286 and GM135218 to M.R.R.) and HHMI. J.L.D. is a Howard Hughes Medical Institute (HHMI) Investigator. N.H.K. was partially supported by Basic Science Research Program through the National Research Foundation of Korea Fellowship funded by the Ministry of Education (2014R1A6A3A03058629). F.E-K was supported University of Tübingen core funding and through the Deutsche Forschungsgemeinschaft (SFB/CRC1101 - project D09) and by a grant from the Reinhard Frank Stiftung (Project ‘helperless plant’) to F.E-K. and J.L.D. L.W. was supported by National key Laboratory of Plant Molecular Genetics, Institute of Plant Physiology and Ecology/Center for Excellence in Molecular Plant Sciences and Chinese Academy of Sciences Strategic Priority Research Program (Type-B; Project number: XDB27040214). We thank Brian Staskawicz and Tiancong Qi for *N. benthamiana nrg adr1* seeds and Keiko Yoshioka, Univ. of Toronto, for *N. benthamiana GCaMP3* seeds. We thank Victoria Madden and Kristen White for handling electron microscopy samples preparation. The Microscopy Services Laboratory, Department of Pathology and Laboratory Medicine, is supported in part by P30 CA016086 Cancer Center Core Support Grant to the UNC Lineberger Comprehensive Cancer Center.

## Author contributions

PJ, NHK, LW and JLD conceived the project. PJ, N-HK, FW, FE-K, WGW, OJF, ADL, SS, KK and LW generated and analyzed data and generated figures. MRR, Z-MP and JLD provided funding and project management. PJ, N-HK, LW and JLD wrote the paper. All other authors contributed edits and comments.

## Materials and Methods

### Recombinant protein cloning, expression, and purification

cDNA of *At*NRG1.1 CC 1-124 (WT, 2K/E and 7K(R)/E) was codon optimized for recombinant protein expression in *E. coli* and subcloned into the pET24a plasmid (Genescript) with a N-terminal (His)6 tag and C-terminal Strep II tag (WSHPQFEK). All proteins were produced in *E. coli* BL21 (DE3) cells using the autoinduction method (*33*) and purified to homogeneity using nickle affinity chromatography followed by size-exclusion chromatography. The selenomethionine (SeMet) incorporated form of 7K(R)/E was produced using SelenoMet^TM^ medium from Molecular Dimensions and 0.1 mM isopropyl-1-thio-D-galactopyranoside (IPTG) for protein induction. Briefly, *E. coli* cells were grown at 37°C until the OD_600_ reached 0.6. The temperature was then decreased to 18°C with protein induction for 18 hours. Cells were pelleted by centrifugation at 6000 x *g* at 4°C for 30 minutes. All cell pellets were resuspended in 5 mL of the lysis buffer (50 mM HEPES pH 8.0, 300 mM NaCl and 2 mM DTT) per gram of cell paste. Phenylmethanesulfonylfluoride (PMSF; 1 mM final concentration), lysozyme, and DNase were added to the cell resuspension immediately prior to lysing. The resuspended cells were lysed using sonication and then centrifuged at 41, 211 x *g* for 20 minutes to remove cell debris. The supernatant was loaded onto a nickel HisTrap 5 mL column (GE Healthcare) pre-equilibrated with 20 mL of the wash buffer (50 mM HEPES pH 8.0, 300 mM NaCl, 30 mM imidazole) at a rate of 3 mL/min. The column was then washed with 100 mL of wash buffer and the bound protein was eluted with elution buffer (50 mM HEPES pH 7.5, 300 mM NaCl, 500 mM imidazole). Elution fractions were monitored via A_280_, elution peak pooled, and further purified using a S200 HiLoad 16/600 column pre-equilibrated with the gel-filtration buffer (10 mM HEPES pH 7.5, 150 mM NaCl and 1mM DTT). The A_280_ elution peak fractions were confirmed by SDS-PAGE, pooled, and the proteins were concentrated to a final concentration of 10 mg/mL. The concentrated proteins were flash-frozen as 30 μL aliquots in liquid nitrogen and stored at -80°C.

### Protein crystallization, data collection, and structure determination

All proteins were crystallized using the sitting drop vapor diffusion technique. A protein solution consisting of ∼10 mg/mL 7K(R)/E native or 7K(R)/E SeMet was mixed at a 1:1 ratio with a precipitant solution composed of 0.1 M MES (pH 5.5) and 1 M potassium sodium tartrate. 7K(R)/E crystals formed within 3 days. Similarly, crystals of 2K/E were produced by mixing a 1:1 ratio of ∼10 mg/mL protein solution with a precipitant solution consisting of 0.1 M HEPES: NaOH (pH 7.5) and 0.3 M magnesium formate. 2K/E crystals formed within 2 days.

Protein crystals of 7K(R)/E native were transferred directly into a mother liquor consisting of the well solution supplemented with ethylene glycol to a final concentration of 20% (v/v). Likewise, 7K(R)/E SeMet and 2K/E crystals were transferred into a mother liquor consisting of the well solution supplemented with glycerol to a final concentration of 20% (v/v). Once transferred, crystals were immediately flash cooled in liquid nitrogen in preparation for x-ray diffraction data collection.

X-ray diffraction data for the 7K(R)/E SeMet construct (M1, M45, M58, and M98) were collected from a single crystal at the 23-IDD beamline using a wavelength tuned to 0.979 Å. Diffraction images were processed with the HKL2000 suite (*34*) and the structure was solved using the single-wavelength anomalous diffraction (SAD) method through the *Autosol* pipeline within the *PHENIX* suite. An initial model was built using the *Autobuild* pipeline within the *PHENIX* suite (*35*). X-ray diffraction data for native crystals of 7K(R)/E were collected at the 23-IDD beamline to a higher resolution than that of the 7K(R)/E SeMet crystals. The native 7K(R)/E diffraction images were processed using the HKL2000 suite and the structure was subsequently solved by molecular replacement using PHASER (*36*) with the 7K(R)/E SeMet build as a search model. X-ray diffraction data for 2K/E were collected at the 23-IDD beamline and the structure was solved by molecular replacement using PHASER with 7K(R)/E as the search model. All models were extended by several rounds of manual model building with COOT (*37*). Successive refinements were performed with Phenix.refine or REFMAC within the CCP4 suite (*35, 38*). Structural analyses were carried out using DALI (*39*) and PyMOL (http://www.pymol.org). Figures were generated using PyMOL. Structures of 7K(R)/E native and 2K/E native are available in the PDB as accessions 7L7V and 7L7W, respectively.

### DNA constructs and vectors used

The *NRG1.1* CDS (wildtype and all analyzed mutants) were synthesized into a pUC57/Kan plasmid (Genescript) including the gateway compatible recombination sites (attR1/attR2). LR reactions (Gateway Cloning Technology, Life Technologies; Carlsbad, USA) were performed to introduce CDS’s into the estradiol-inducible destination vector pMDC7 with a C-terminal YFP-HA CDS (*40*), or the 35s-driven destination vector pGWB641 (*41*). The clones of NRG1.1 1- 180, ΔN16 and structure-derived mutants (all C-terminal GFP-fusions) were constructed in pICSL86922 (35S promoter and TMV omega enhancer) using Golden Gate cloning (*42*).

### Bacterial strains and growth conditions

*Agrobacterium tumefaciens* strain GV3101/pMP90 were grown in LB media at 28**°C**. Antibiotic concentrations used (in g/mL) Kanamycin 100, Gentamycin 50 and Spectinomycin 100, Rifampicin 100. Constructs were transformed into *Agrobacterium tumefaciens* strain GV3101/pMP90 and used for transient expression in *Nicotiana benthamiana*. NRG1.1 WT and variants were transiently expressed in *Nicotiana benthamiana*.

### Determination of NRG1.1 localization using confocal imaging

Briefly, *Agrobacterium tumefaciens* strains were grown overnight at 28°C in LB media containing the appropriate antibiotics. The overnight cultures were centrifuged for 8 min at 8,500 rpm and the pellets were resuspended in induction buffer (10 mM MgCl_2_, 10 mM MES pH 5.6, 150 µM acetosyringone). The OD_600_ was adjusted to 0.05 (35S::P19) and 0.3 for NRG1.1 constructs and the specific subcellular marker constructs (35S::BRI1-mRFP – a gift from Klaus Harter, UBQ10::VMA12-RFP – a gift from Karen Schuhmacher – and 35S::PLC2-CFP). Samples were mixed as indicated. Agrobacteria mixtures were infiltrated into young leaves of 4-6 weeks old *N. benthamiana* WT plants using a 1-ml needleless syringe. The *N. benthamiana* plants were grown on soil under 12h light / 12h dark cycles (24°C/22°C, 65% humidity). Induction was done 24 hours post infiltration using either 20 µM β-estradiol (Sigma-Aldrich; St. Louis, USA) and 0.001% [v/v] Silwet L-77 by spraying. Leaves were imaged for protein localization between 6 h- 24 h post induction with the confocal laser scanning microscope LSM880 from Zeiss (Oberkochen, Germany), using C-apochromat 40x or 63x/1.2 W Korr FCS M27 water-immersion objectives and the ZENblack software. YFP was excited using a 514 nm laser collecting emission between 516-556 nm; RFP was excited using a 561 nm laser with an emission spectrum of 597-633 nm, CFP was excited with a 458 nm laser and the emission spectrum was 463-513 nm. Focal plane images were processed with the ZENblue software (Zeiss) for adjustment of brightness and contrast. Maximum Z-projection images were processed with ImageJ/Fiji (*43*).

### UV-light imaging of cell death phenotypes

Briefly, *A. tumefaciens* strains were infiltrated into *N. benthamiana* leaves as described above. The OD_600_ was adjusted to 0.1 (35S::P19) and 0.8 for NRG1.1 and ADR1 constructs. For Ca^2+^ inhibitor treatments, 2 mM LaCl_3_ or GdCl_3_ was added to the infiltration media prior to infiltration. Induction was done 24 hours post infiltration using 100 µM β-estradiol (Sigma-Aldrich; St. Louis, USA) and 0.002% [v/v] Silwet L-77 by spraying. 1-2 days post induction the leaves were placed under UV lamps (B-100AP, UVP) and photographed using a digital camera with a yellow filter.

### In planta semi-quantification of Ca^2+^ influx

Briefly, *A. tumefaciens* strains were infiltrated into transgenic GCaMP3 *N. benthamiana (44)* leaves and protein expression induced as described above. Leaf discs (0.5 cm diameter) were placed on 200 uL ddH_2_O in Nunclon 96 Flat Bottom Black plates (Thermo-Fisher) and equilibrated for 1 h in room temperature. After equilibration, GCaMP3 fluorescence was recorded using TECAN Infinite M200 Pro plate-reader, using excitation at 470 nm (7 nm bandwidth) and emission detection at 525nm (20nm bandwidth) with 20 µs integration time and 5 ms settle time. Absolute fluorescence values for each experiment were normalized to the untreated control value as F/F_UT_ (where F was the measured fluorescence at a given time point and F_UT_ was the averaged measurement for water-equilibrated uninfiltrated samples at each time points). Plotted values are averages of six replicate leaf discs, with the SE represented by error bars.

### Ion leakage measurements

Briefly, *A. tumefaciens* strains were infiltrated into *N. benthamiana* leaves and protein expression induced as described above. 12 leaf discs (0.5 cm diameter) from 3 independent plants were collected into clear tubes with 20 ml of distilled water and incubated at room temperature under continuous light (three replicates per sample). Changes in conductivity were measured at the indicated time points with an Orion Model 130 (Thermo-Fisher).

### HeLa cell transfection

Human cervical cancer cells (HeLa) were cultivated in DMEM medium containing 10% fetal bovine serum and 1% penicillin / streptomycin in a CO_2_ incubator at 37C. Cells were grown in 12- well plates overnight to 70-90% confluence and co-transfected with the “sleeping beauty” pCMV(CAT)T7-SB100X vector containing the sleeping beauty transposase and the pSBtet-Pur vectors - a gift from Long Ping V. Tse and Bill Marzluff - containing the synthesized, human codon optimized, NRG1.1 constructs (Genescript) at a 1:10 ratio respectively. GASDERMIN-D L192D N-terminal pore-forming domain (aa 1-275), a potent inducer of cell death, and the loss of function GASDERMIN-D L192D DN5 were used as controls (*45-47*). All constructs were YFP- tagged at the C-terminus. Briefly, for each co-transfection, 25μL of Opti-MEM medium was mixed with 0.75uL of Lipofectamine 3000 (Invitrogen) in one tube while 1μg of the plasmid mix was added to 25μL of Opti-MEM medium and 1uL of P3000 reagent in another tube. Solutions were mixed together and incubated at room temperature for 15 minutes before being added to the cell culture. After 2 days, puromycin 1μg.mL^-1^ (Sigma) was added to the cultures to select transfected cells.

### Protein Assays

*Agrobacterium tumefaciens* strains were infiltrated into *N. benthamiana* leaves as described above. The OD_600_ was adjusted to 0.1 (35S::P19) and 0.8 for NRG1.1 constructs. Induction was done 24 hours post infiltration using 100 µM β-estradiol (Sigma-Aldrich; St. Louis, USA) and 0.002% [v/v] Silwet L-77 by spraying. Tissues for protein assays were collected 6 h post induction.

Total protein extract was prepared by adding total protein buffer [100 mM Tris-Cl pH 7.5, 150 mM NaCl, 1% Triton X-100, 1 mM EDTA pH 8.0, 0.1% SDS, 10 mM dithiothreitol (DTT) and 1x Sigma plant protease inhibitor cocktail] to the homogenized tissue at a ratio of 5 uL per mg (FW) tissue. Lysate was cleared by centrifuge at 20,000 x *g* for 30 min at 4°C.

For microsomal fractionation, sucrose buffer [20mM Tris (pH 8), 0.33M sucrose, 1mM EDTA, 5mM DTT and 1x Sigma plant protease inhibitor cocktail] was added to the homogenized tissue at a ratio of 5 uL per mg (FW) tissue. The extract was centrifuged at 2,000 x *g* for 5 minutes at 4°C to remove debris. The resulting supernatant was transferred to a new tube and designated as total protein. Cytoplasmic fraction was prepared by harvesting the supernatant after spinning the total protein fraction at 20,000 x *g* for 1 h at 4°C. The total membrane fraction was prepared from the resulting pellet by resuspending in 200 uL of buffer B (Minute^TM^ plasma membrane protein isolation kit, Invent Biotechnologies) and spinning at 7,800 x *g* for 15 min at 4°C. The pellet was designated as total membrane fraction. The supernatant was transferred to 2 mL Eppendorf tube and mixed with 1.6 mL cold PBS buffer mixed by vortexing and spun at 16,000 x *g* for 45 min at 4°C to pellet the plasma membrane fraction.

For blue native polyacrylamide gel electrophoresis, modified GTEN buffer (*48*) [10% glycerol, 100 mM Tris-Cl pH7.5, 1mM EDTA, 150mM NaCl, 5mM DTT, 0.5% n-dodecyl-B-D-maltoside (DDM) and 1x Sigma plant protease inhibitor cocktail] was added to the homogenized tissue at a ratio of 5 uL per mg (FW) tissue. Lysate was cleared by centrifuge at 20,000 x *g* for 30 min at 4°C. The supernatant was transferred to a new tube and mixed with 4x NativePAGE Sample buffer (Invitrogen).

For SDS-PAGE, proteins were mixed with 4x SDS-PAGE loading buffer and heated for 5 min at 95°C and resolved in 10% SDS-PAGE gels. Separated proteins were transferred to nitrocellulose membrane (GE Healthcare). For BN-PAGE, proteins were resolved in 3-12% and NativePAGE^TM^ (Invitrogen) gels, respectively, then transferred to PVDF membrane using iBlotTM 2 gel transfer device according to the instructions from the manufacturer. Membranes were blocked with 5% non-fat milk TBST and probed with primary anti-GFP (Roche, 1:1000) and secondary HRP-conjugated anti-mouse antibody (R&D Systems, 1:5000).

### Cell death observation in HeLa cells

After 2 passages, cells were seeded on 8 well chamber slides (Nunc Lab-Tek 155411), and protein expression was induced by addition of doxycycline 1μg.mL^-1^ (PHR1145, Sigma). 6 hours after doxycycline addition, the cells incubated in incubation buffer (10 mM HEPES, 140 mM NaCl, and 2.5 mM CaCl_2_, pH 7.4) containing 1μg.mL^-1^ propidium iodide (PI) for 5 min and observed under a confocal microscope LSM710 AxioObserver (Zeiss), using the Plan-Apochromat 10X/0.45 M27 objective. YFP was excited using a 514 nm laser collecting emission between 520-559 nm; PI was excited using a 561 nm laser with an emission spectrum of 590-627 nm. Percentage of cell death was determined as the number of PI-positive cells amongst YFP positive cells.

### Plasma membrane pore quantification with scanning electron microscopy

Transfected cells were seeded on glass 12mm glass slides and grown overnight to 70-90% confluence. 6 hours after doxycycline treatment, the media was removed and cells were washed in PBS and fixed in 2.5% glutaraldehyde in 0.1M sodium cacodylate at room temperature for 30 minutes and stored for several days at 4°C. To better preserve the membrane structure, decrease shrinkage artefacts and reduce the metal coating thickness, the cells were post- fixed in 1% osmium tetroxide in 0.1M sodium cacodylate buffer for 15 minutes followed by subsequent treatment with 2% tannic acid in water for 15 minutes and 1% osmium tetroxide in water for 15 minutes (*49*). The coverslips were rinsed in deionized water prior to gradual dehydration with ethanol (30%, 50%, 75%, 100%, 100%) and critical point dried using carbon dioxide as the transitional solvent (Samdri-795 critical point dryer, Tousimis Research Corporation, Rockville, MD). Coverslips were mounted on 13mm aluminum stubs using double-sided carbon adhesive and sputter coated with 3 nm of gold-palladium alloy (60 Au:40 Pd, Cressington Sputter Coater 208HR, model 8000-220, Ted Pella, Redding, CA). Images were taken using a Zeiss Supra 25 FESEM operating at 5 kV, using the InLens detector, 30µm aperture, and ∼5 mm working distance (Carl Zeiss SMT Inc., Peabody, MA). Plasma membrane pores were observed at 200,000 X magnification. Two micrographs were taken per cell and at least 11 cells were observed per cell line. Experiment was performed twice with similar results.

### Imaging of [Ca^2+^]_i_ in HeLa cells

Transfected HeLa cells were seeded on eight-well chambered coverglasses (Nunc Lab-tek 155411) and grown overnight in a CO_2_ incubator at 37°C. 6 to 8h post induction with 1 μg mL^-1^ doxycycline, a Fura-2-based Ca^2+^ imaging assay was performed as previously described (*50*). Cells were loaded with the Ca^2+^ sensitive dye Fura-2AM (5 µM; Sigma) and incubated in a low [CaCl_2_] standard buffer containing 130 mM NaCl, 3 mM KCl, 0.6 mM MgCl_2_, 10 mM glucose, 10 mM HEPES, pH7.4 (adjusted with NaOH), and 0.1 mM CaCl_2_ for 30 min. A 7.5 mM CaCl_2_ standard buffer was added with a peristaltic pump (Dynamax RP-1, Rainin) to adjust final [CaCl_2_] to 2.5 mM. Fura-2 fluorescence imaging was performed using the Zeiss Axiovert 200 microscope equipped with two filter wheels (Lambda 10-3; Sutter Instruments) and a CMOS camera (Prime sCMOS; Roper Scientific). Excitation was at 340 nm and 380 nm, and emission images at 510 nm were collected using the MetaFluor Fluorescence Ratio Imaging Software (Molecular Devices). Ratiometric (*F*340 nm/*F*380 nm) image data from 10 cells per well were analyzed. Experiments were carried out at room temperature (22–24°C). Delta ratio was found by calculating the difference of the average fluorescence ratio from 10 cells from 6 time points after switching to high [CaCl_2_] and those same cells at 6 time points before switching to high [CaCl_2_].

### Imaging of [Cl^-^]_i_ in HeLa cells

Transfected HeLa cells were seeded on eight-well chambered borosilicate cover glasses (Nunc Lab-tek 155411) and grown overnight. At 6 to 8h post induction with 1 μg mL^-1^ doxycycline, cells were subjected to a 6-Methoxy-N-ethylquinolinium iodide (MEQ) -based Cl^-^ assay as previously described (*51*). DihydroMEQ was prepared freshly from MEQ. 5 mg of MEQ were dissolved in 100 μL of distilled water. The MEQ solution was flushed with nitrogen and reduced to dihydroMEQ by addition of 32 μmol of sodium borohydride with continuous purging with nitrogen. DihydoMEQ was extracted with chloroform, dried with anhydrous sodium sulfate and the organic extracts were evaporated under a vacuum. DihydroMEQ was dissolved in DMSO and added to a low chloride buffer (5 mM KCl, 20 mM HEPES, 90 mM mannitol) at a final concentration of approximately 50 μM. Cells were incubated for 5-10 minutes in the low chloride, dihydroMEQ containing buffer, rinsed and incubated 15 minutes in 200 μL of low chloride buffer before confocal observation. Basal MEQ fluorescence was imaged for 5 min before addition of 100 μL of 200 mM NaCl. Data was analyzed using FiJi (*43*). The MEQ fluorescence of 12 cells was followed up in each experiment. Experiments were carried out at room temperature.

---

**Fig. S1.**
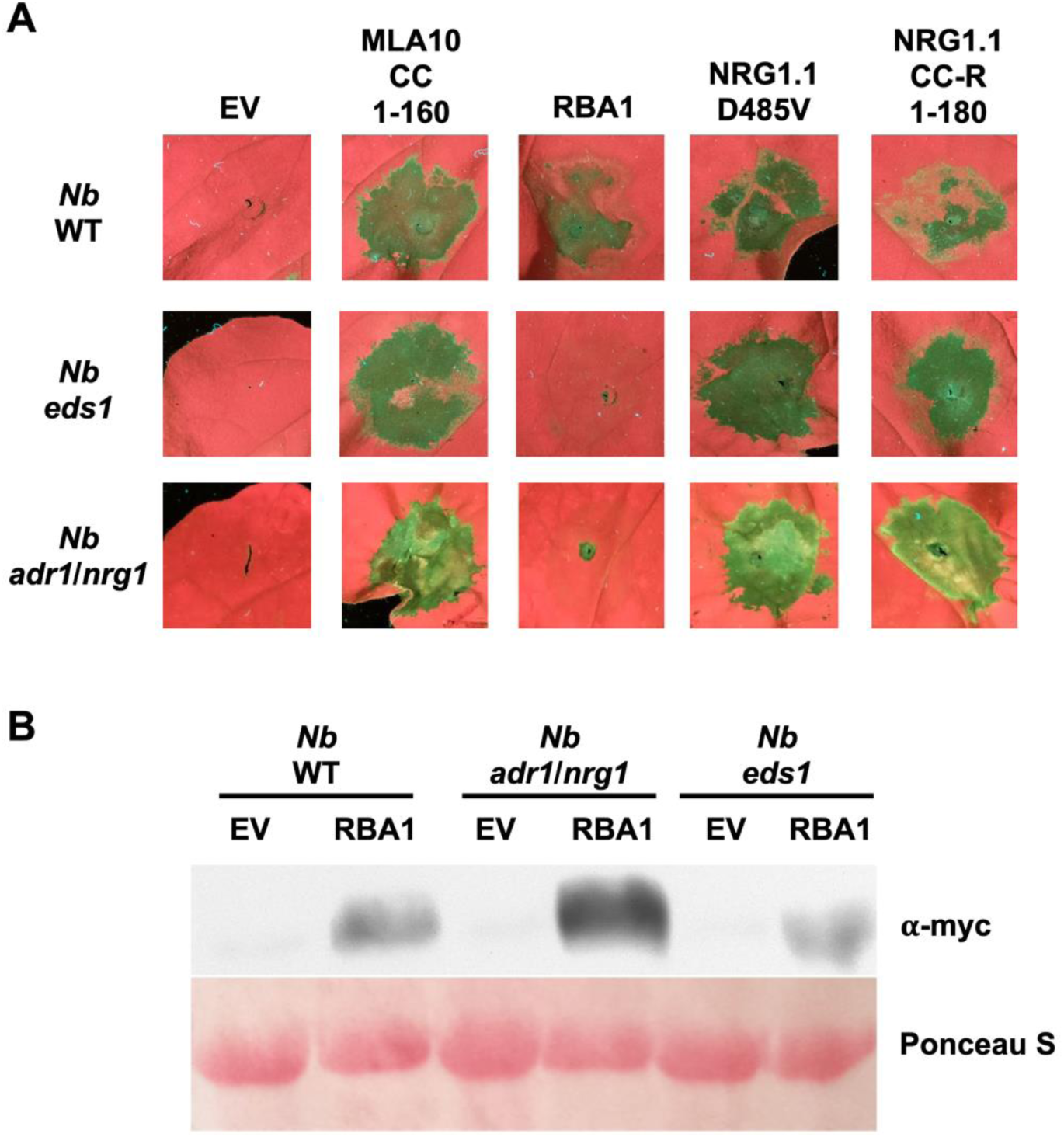
Auto-active AtNRG1.1 D485V triggers cell death independently of EDS1 and NRG1/ADR1 in *Nb*. (**A**) UV-image of *Nicotiana benthamiana (Nb)* leaves of indicated genotypes *Nb*, *Nb eds1* or *Nb adr1 nrg1* infiltrated with Agrobacterium harboring indicated Myc-tagged constructs to observe cell death phenotype 3 days post infiltration. A 180 aa N-terminal fragment of NRG1.1 is a control for cell death inductions (*52*). (**B**) Myc-tagged RBA1 proteins extracted from harvested leaf tissues 26 hours post infiltration were resolved by SDS-PAGE and blotted for Myc.

**Fig. S2.**
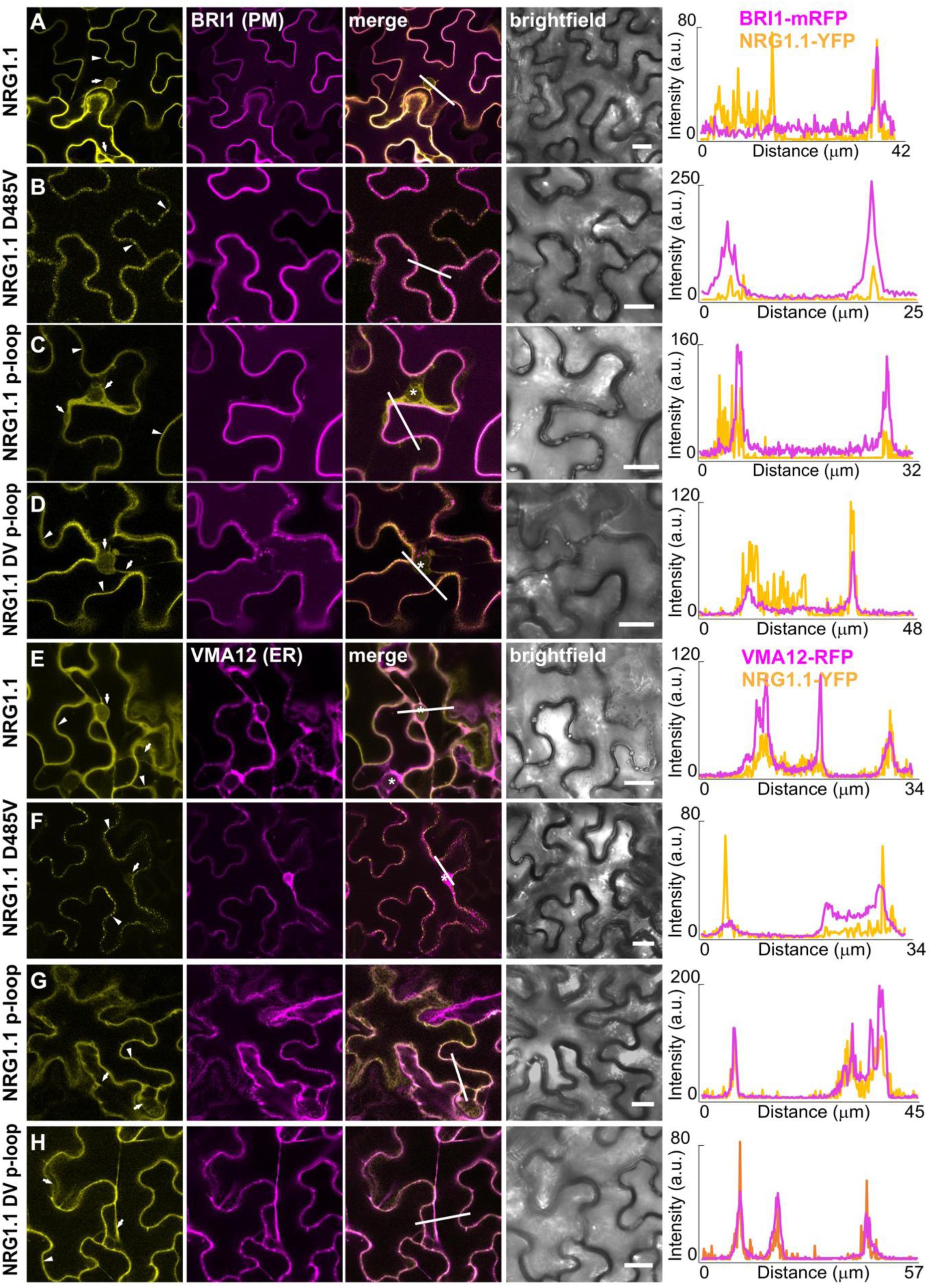
Active NRG1.1 localizes to the plasma membrane. **A** to **D** are representative images showing the (co-)localization of wildtype NRG1.1 (**A**), NRG1.1 D485V (**B**), NRG1.1 p-loop (**C**) and NRG1.1 DV p-loop (**D**) with the plasma membrane (PM) localized BRI1-mRFP. **E** to **H** are representative images showing the (co-) localization of wildtype NRG1.1 (**E**), NRG1.1 D485V (**F**), NRG1.1 p-loop (**G**) and NRG1.1 DV p-loop (**H**) with the endoplasmic reticulum (ER) localized VMA12-RFP. Notably, active NRG1.1 D485V colocalizes poorly with VMA12 (ER) and strongly with BRI1 (PM) whereas the WT, inactive p-loop and DV p-loop colocalize strongly with the ER marker VMA12 and partially with the PM marker BRI1. Indicated proteins were transiently expressed in 4-6 week-old wildtype *Nicotiana benthamiana* plants. Expression of the indicated NRG1.1 constructs was induced 24 hours post infiltration with 20uM β-estradiol + 0.001% [v/v] Silwet L-77 and confocal imaging was performed 6 to 24 hours post induction. Arrowheads in **A** to **H** point to plasma membrane and/or punctate structure localization and arrows indicate endoplasmic reticulum localization. Lines in the merged images indicate the path of the intensity profile shown on the right and are pointing from left to right or up to down. Scale bars = 20 μm.

**Fig. S3.**
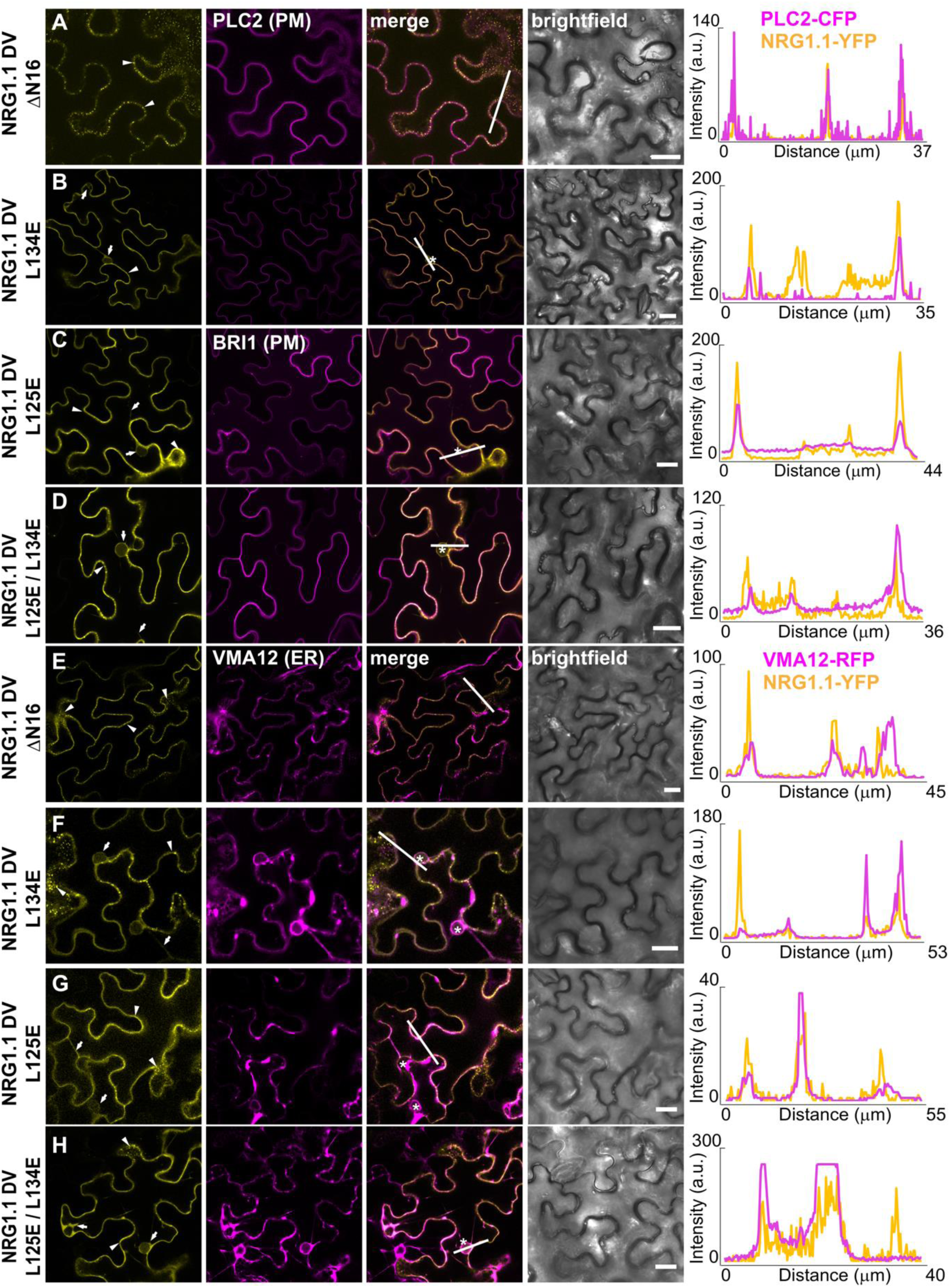
Impact of structure-derived mutations on NRG1.1 localization. **A** to **D** are representative images showing the (co-) localization of NRG1.1 DV ΔN16 (**A**), NRG1.1 DV L134E (**B**), NRG1.1 DV L125E (**C**) and NRG1.1 DV L125E/L134E (**D**) with the plasma membrane (PM) localized BRI1-mRFP or PLC2-CFP, as indicated. **E** to **H** are representative images showing the (co-)localization of NRG1.1 DV ΔN16 (**E**), NRG1.1 DV L134E (**F**), NRG1.1 DV L125E (**G**) and NRG1.1 DV L125E/L134E (**H**) with the endoplasmic reticulum (ER) localized AtVMA12-RFP. Notably, NRG1.1 DV ΔN16 and NRG1.1 DV L125E colocalize strongly with the PM marker PLC2 or BRI1 and less so with the ER marker VMA12, whereas NRG1.1 DV L134E and NRG1.1 DV L134E/L125E colocalize strongly with the ER marker VMA12 and less so with the PM markers PLC2 or BRI1. Indicated proteins were transiently expressed in 4-6 week-old wildtype *Nicotiana benthamiana* plants. Expression of indicated NRG1.1 constructs was induced 24 hours post infiltration with 20uM β-estradiol + 0.001% [v/v] Silwet L-77 and confocal imaging was performed 6 to 24 hours post induction. Arrowheads in **A** to **H** point to plasma membrane and/or punctate structure localization and arrows indicate endoplasmic reticulum localization. Lines in the merged images indicate the path of the intensity profile shown on the right and are pointing from left to right or up to down. Scale bars = 20 μm.

**Fig. S4.**
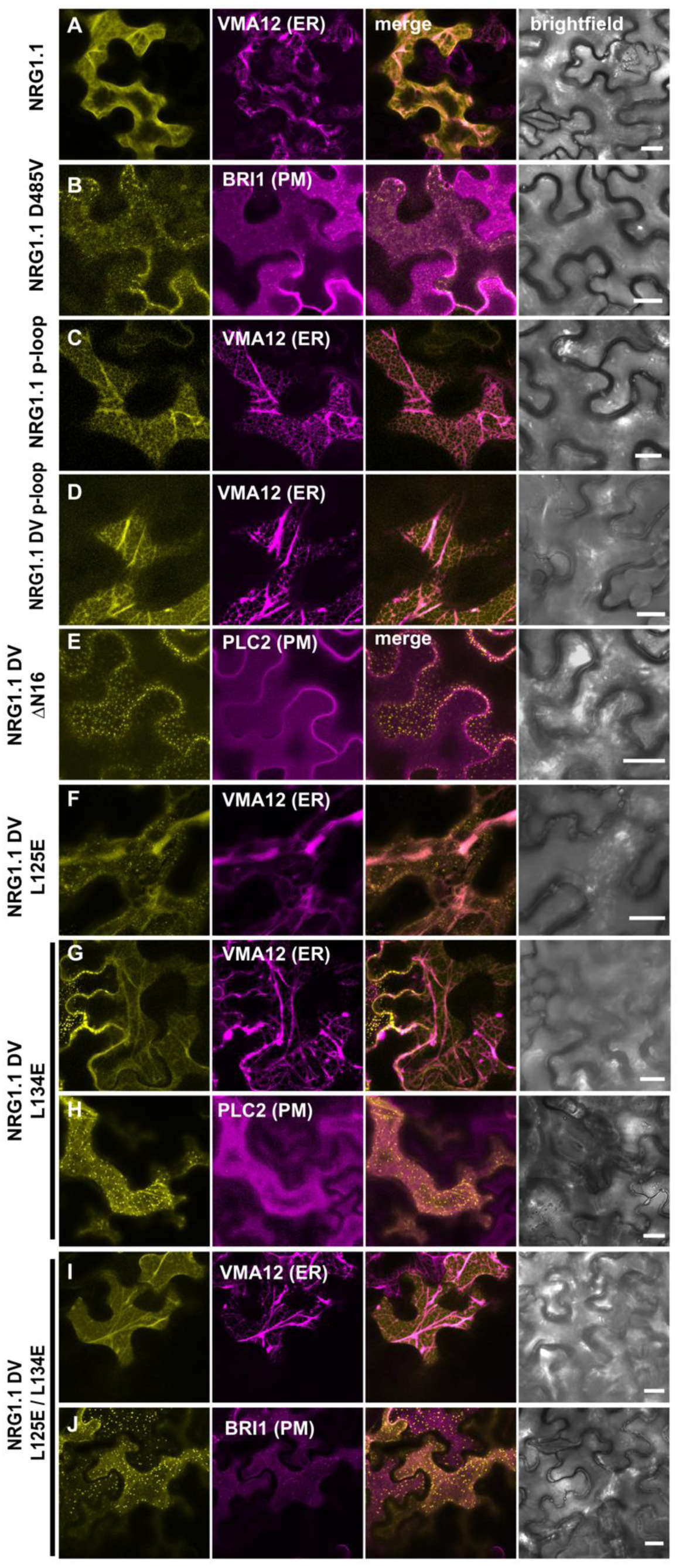
Surface view of NRG1.1 WT and variants. **A** to **J** are representative surface view images showing the (co-) localization of wildtype NRG1.1 (**A**), NRG1.1 D485V (**B**), NRG1.1 p-loop (**C**) and NRG1.1 DV p-loop (**D**), NRG1.1 DV ΔN16 (**E**), NRG1.1 DV L125E (**F**), NRG1.1 DV L134E (**G, H**) and NRG1.1 DV L125E/L134E (**I,J**) with either the plasma membrane (PM) localized AtBRI1-mRFP or AtPLC2-CFP or the endoplasmic reticulum (ER) localized AtVMA12-RFP. Importantly, PM localization is not uniform. PM localized NRG1.1 DV, NRG1.1 DV ΔN16 and NRG1.1 DV L134E were found in puncta at the PM whereas inactive NRG1.1 WT, p-loop and DV p-loop were associated with characteristic ER strands In some examples, NRG1.1 L134E and NRG1.1 DV L134E/L125E, to a much lesser extend NRG1.1 L125E, exhibited puncta at the PM which may reflect some residual PM localization (see also Fig 2C). Indicated proteins were transiently expressed in 4-6 week-old wildtype *Nicotiana benthamiana* plants. Expression of indicated NRG1.1 constructs was induced 24 hours post infiltration with 20uM β-estradiol + 0.001% [v/v] Silwet L-77 and confocal imaging was performed 6 to 24 hours post induction. Surface view images were constructed with the ImageJ software by combining 7-15 Z-stacks of the surface of individual cells into a maximum projection image. Note: For both the NRG1.1 DV L134E and NRG1.1 DV L125E/L134E mutants, we observed cells showing both strong ER localization (**G** and **I**) and the punctate localization pattern at the PM (**H** and **J**). Scale bars = 20 μm.

**Fig. S5.**
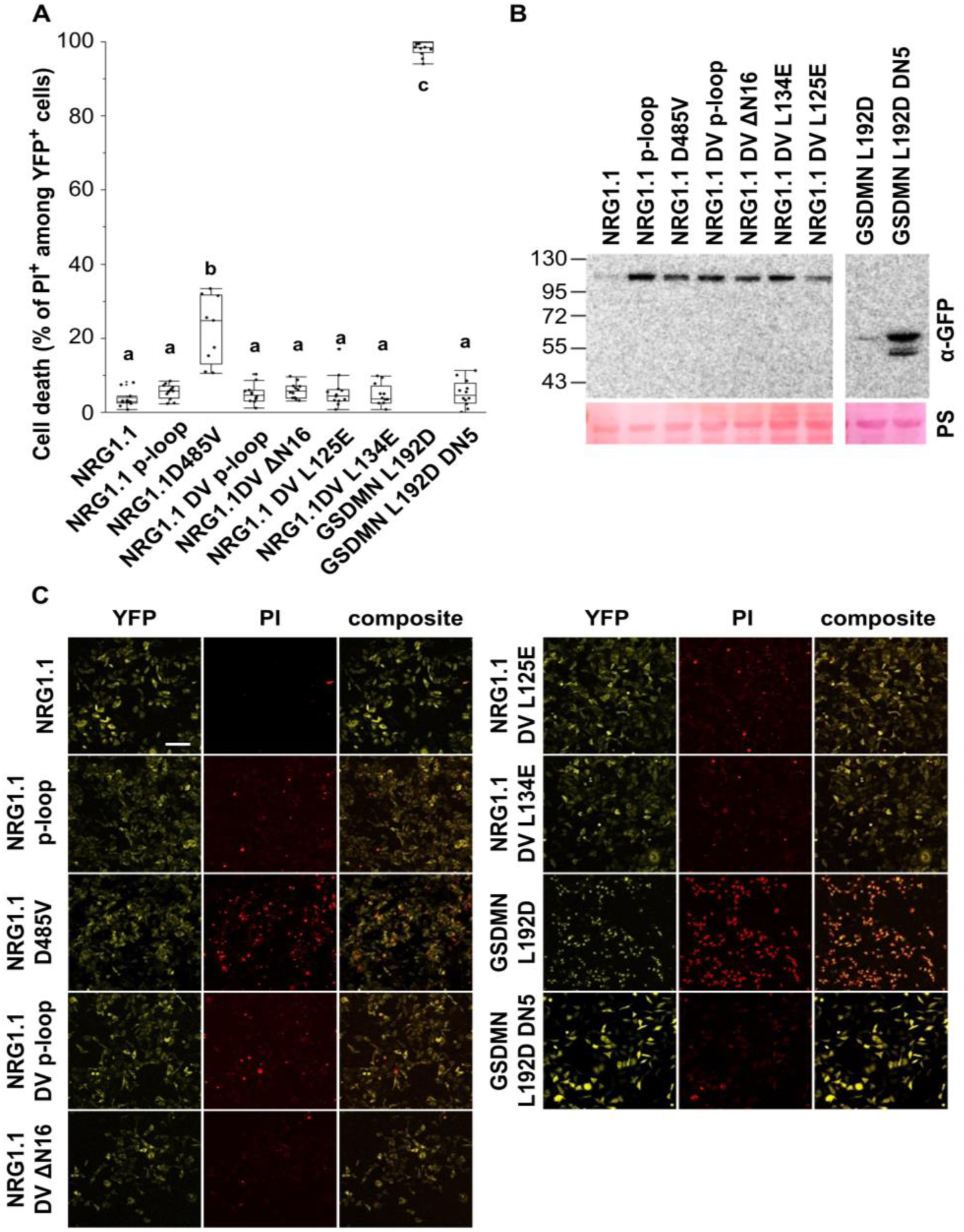
NRG1.1 cell death induction function is conserved in HeLa cells. **A** Cell death percentage observed 6 hours post-induction (hpi) of protein expression with 1μg.ml^-1^ doxycycline. Cells were stained with propidium iodide (PI) and observed with a confocal microscope. The cell death is calculated as the percentage of PI^+^ cells among the YFP^+^ cells. Each data point is the average of counts from at least 3 different confocal images. The data presented here summarize 3 independent experiments. Letters represent statistical difference (ANOVA with post-hoc Tukey HSD test, p-value<0.0001). NRG1.1 cell death function can be recapitulated in HeLa cells. **B** Western blot confirming protein accumulation at 6hpi with doxycycline. Ponceau S stands for Ponceau Staining. **C** Representative confocal fluorescence images used for cell death quantification in **A**. Scale bar is 100μm.

**Fig. S6.**
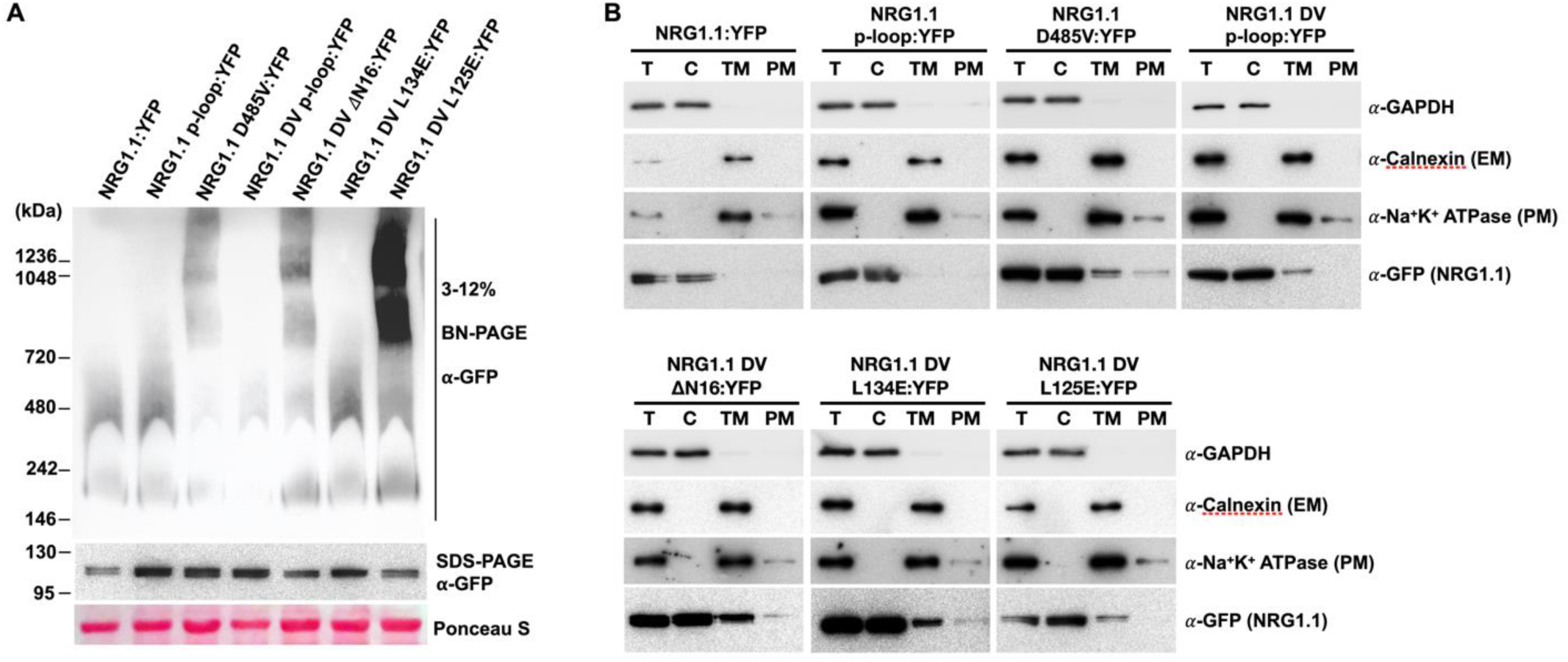
Active NRG1.1 oligomerizes and localizes to the plasma membrane in HeLa cells. **A** NRG1.1 DV forms high molecular weight oligomers in HeLa cells. Duplicate samples were analyzed by either SDS-PAGE (monomers) or BN-PAGE (oligomers). YFP tagged proteins were detected with anti-GFP antibody. Ponceau S stain is shown as loading control. **B** NRG1.1 localizes to the plasma membrane in HeLa cells. HeLa cells expressing indicated constructs were fractionated to Total proteins (T), cytosolic (C), total membrane (TM) and plasma membrane (PM) fractions and verified by marker proteins: Cytosol, GAPDH; endoplasmic reticulum membrane (EM), Calnexin; PM, Na^+^K^+^ATPase. NRG1.1 proteins were detected by anti-GFP antibody.

**Fig. S7.**
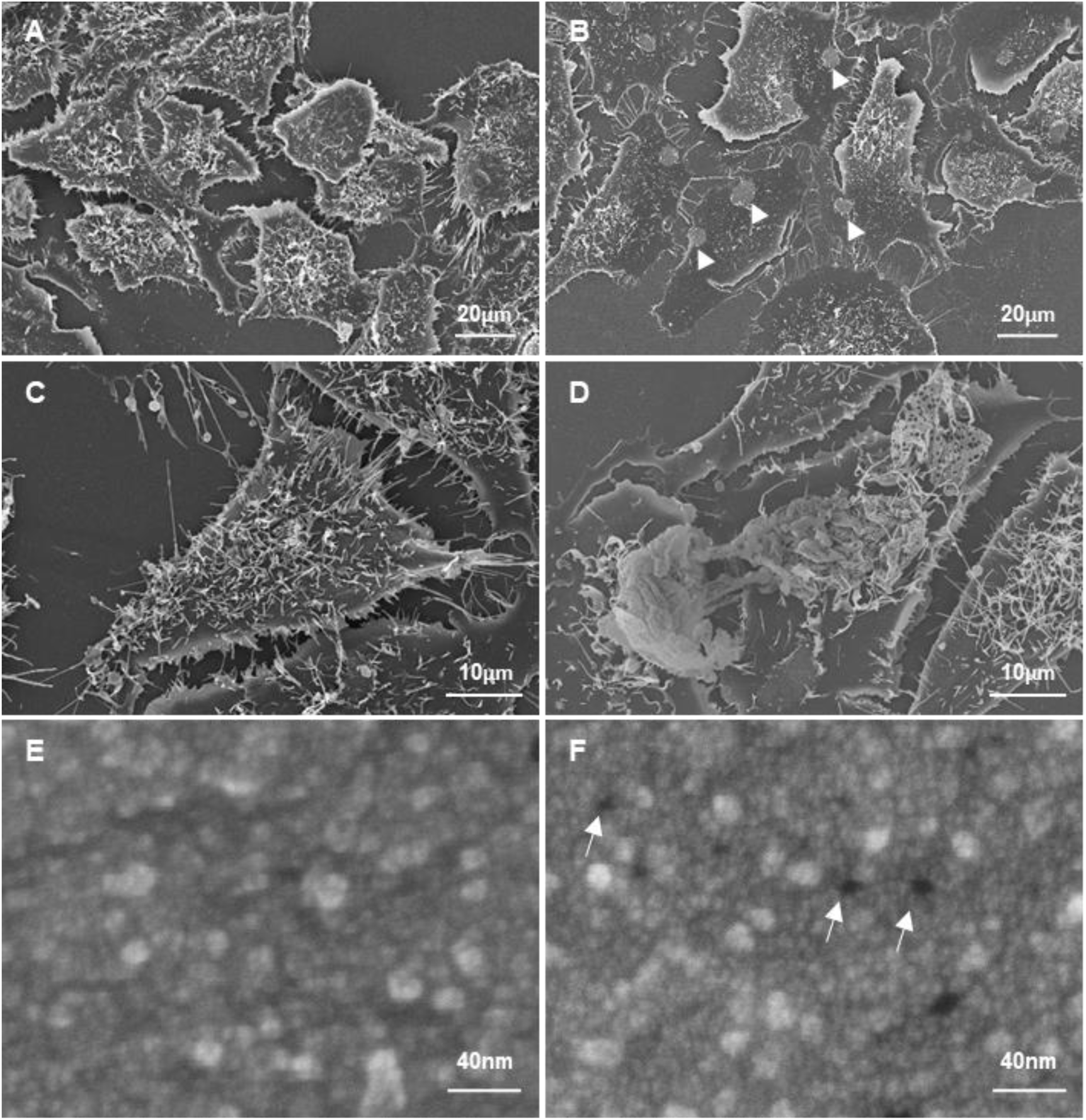
Morphology of NRG1.1 D485V expressing cells. Representative scanning electron microscopy pictures of HeLa cells 6 hours post doxycycline induction of inactive NRG1.1 D485V p-loop (**A**, **C** and **E**) or active NRG1.1 D485V (**B**, **D** and **F**). HeLa cells expressing the active NRG1.1 D485V (**B**) seem to have lost plasma membrane integrity and consequently appear flat with fewer microvilli compared to cells expressing the inactive NRG1.1 D485V p-loop (**A**). White arrowheads indicate large holes in the plasma membrane visible in many NRG1.1 D485V expressing cells. NRG1.1 D485V expressing cells occasionally showed “bubbling” reminiscent of pyroptosis (**D**). Expression of active NRG1.1 D485V was associated with a large number of plasma membrane pores (**F**) compared to the inactive control (**E**). White arrows indicate some plasma membrane pores.

**Fig. S8.**
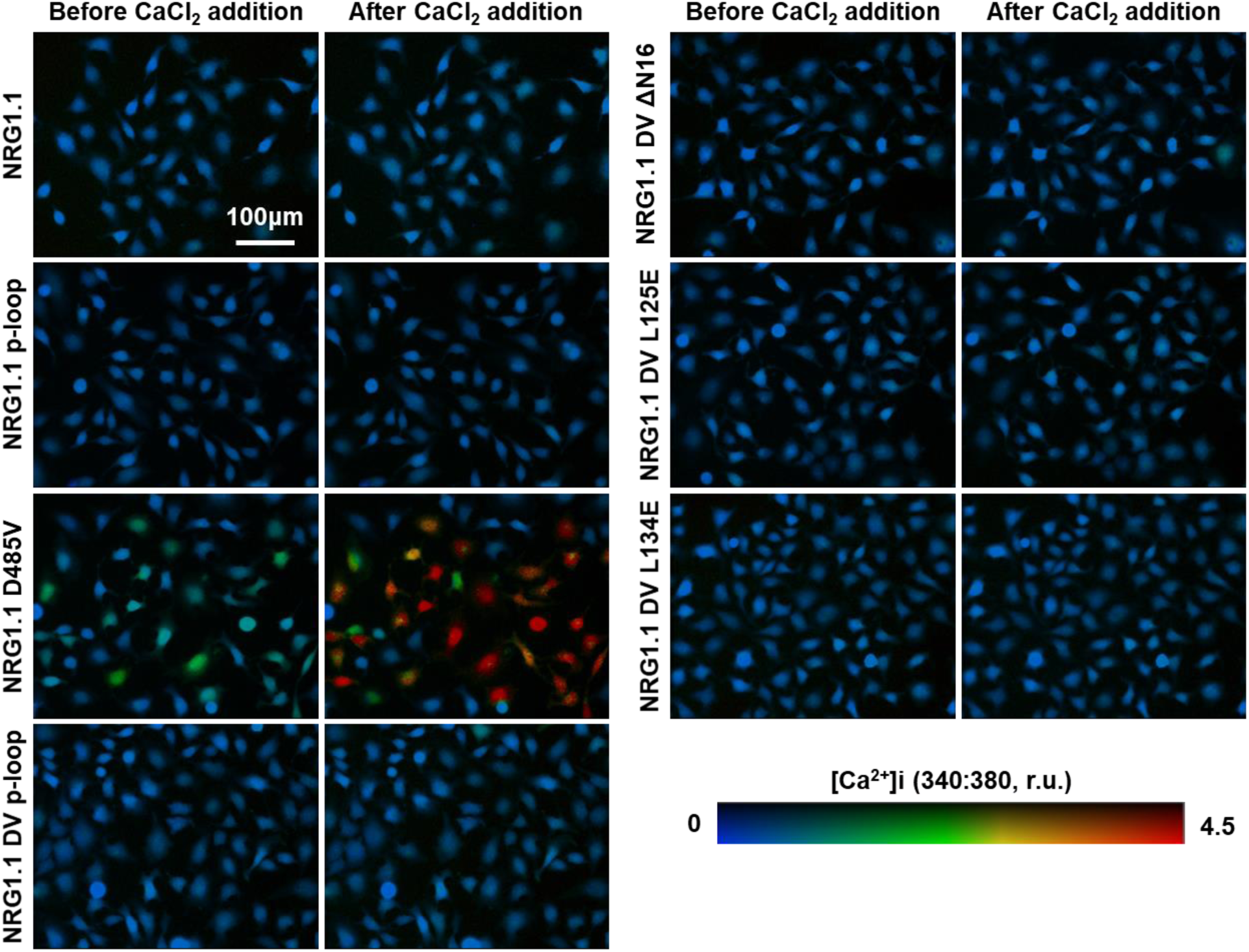
Fura-2-based quantification of intracellular [Ca^2+^] in HeLa cells expressing NRG1.1 and variants. Representative fluorescence microscopy images of HeLa cells expressing NRG1.1 WT or variant, as indicated, before or 2 minutes after 2.5mM CaCl_2_ addition. Intracellular [Ca^2+^] was calculated from Fura-2 emission ratios (340:380 nm) and scaled using a pseudo-color bar. The ratios were quantified in Fig 3K.

**Fig. S9.**
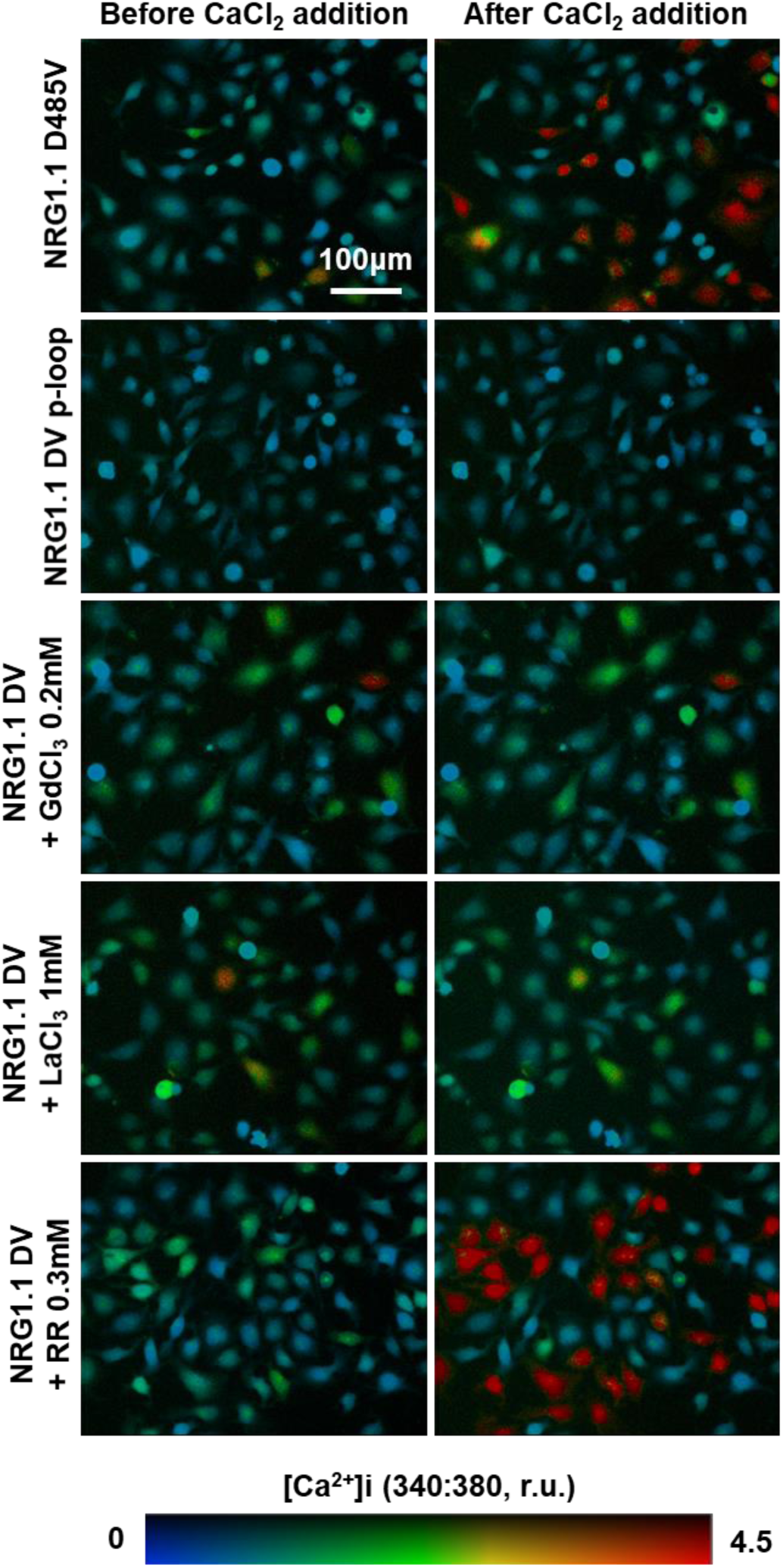
Fura-2-based quantification of Calcium channel blockers impact on intracellular [Ca^2+^] in NRG1.1 D485V-expressing HeLa cells. Representative fluorescence microscopy images of HeLa cells expressing active NRG1.1 D485V in the presence of typical calcium channel blockers, before or 2 minutes after 2.5 mM CaCl_2_ addition. Intracellular [Ca^2+^] was calculated from Fura-2 emission ratios (340:380 nm) and scaled using a pseudo-color bar. The ratios were quantified in Fig3 L.

**Fig. S10.**
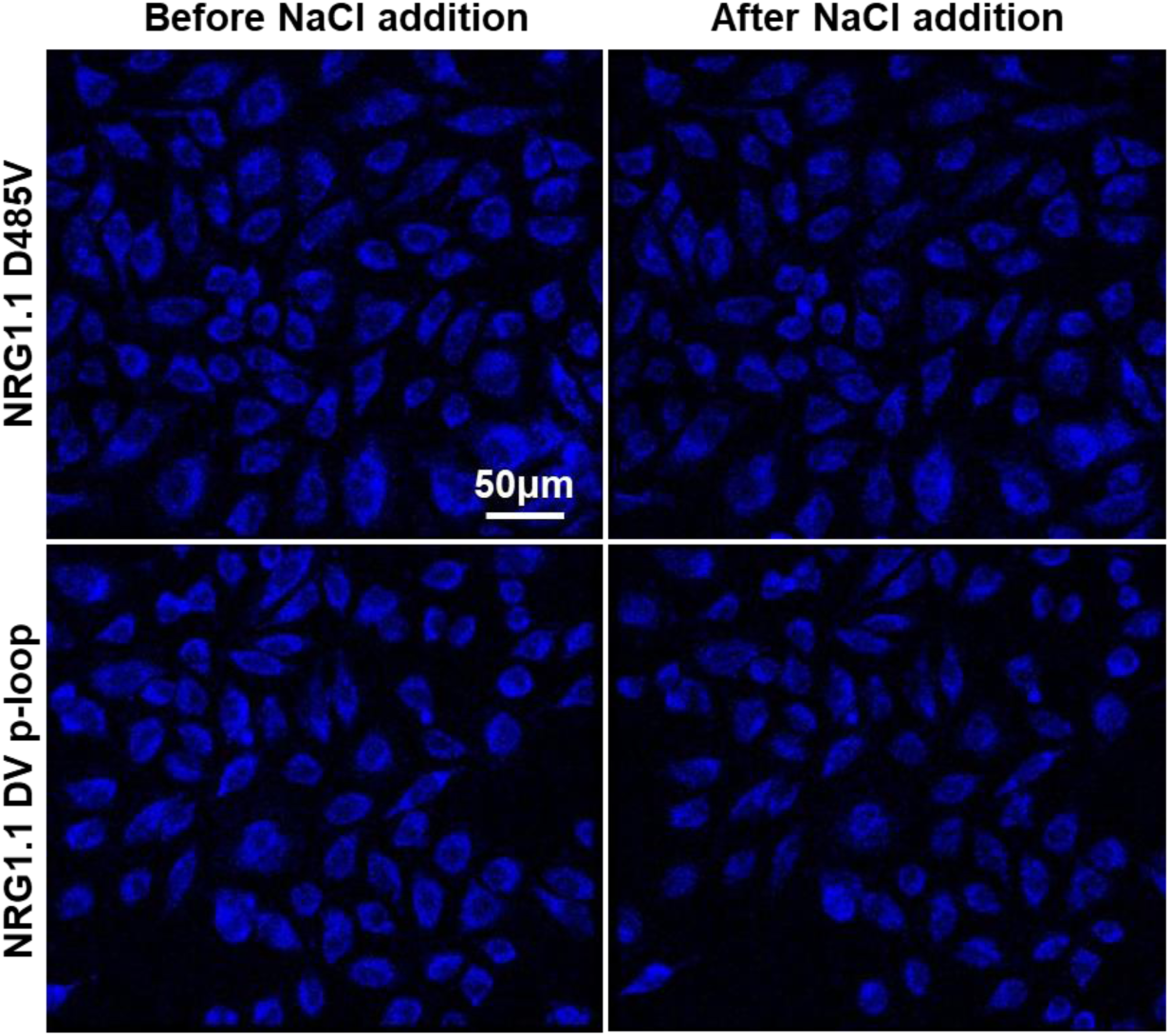
MEQ-based quantification of intracellular [Cl^-^] in NRG1.1 D485V and NRG1.1 DV p-loop-expressing HeLa cells. Representative fluorescence microscopy images of HeLa cells expressing active NRG1.1 D485V or suppressed NRG1.1 DV p-loop, before or 2 minutes after 200 mM NaCl addition. Intracellular [Cl^-^] was visualized from MEQ fluorescence quenching (emission at 440nm). The signal was quantified in Fig3 M.

**Fig. S11.**
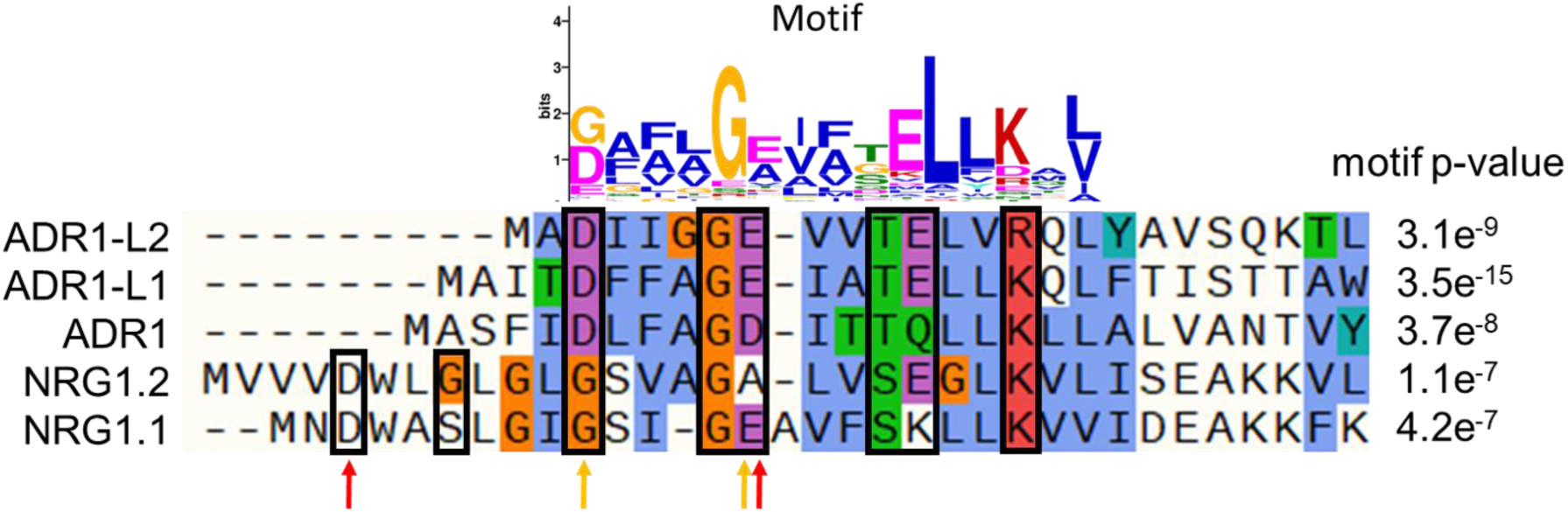
The RNL N-terminal motif aligned to Arabidopsis ADR1s and NRG1s N-termini. *De novo* motif discovery algorithm (MEME, meme-suite.org, (*53*)) run on 334 RNL N-terminal sequences (pfam.xfam.org; Table S3) allowed the identification of a conserved pattern of charged, polar or glycines residues (black boxes) separated by 2 to 3 hydrophobic residues (Motif E-value = 2.8e-1139, 331 occurrences). The motif is partially degenerated in NRG1.1 and NRG1.2 with K19 (NRG1.1) and A17 (NRG1.2) replacing E residues. Interestingly, NRG1.1 and NRG1.2 possess an N-terminal extension comprising D-G/S residues that seem to follow the same spacing pattern. Red and orange arrows indicate the residues that were mutated in NRG1.1 and ADR1 respectively.

**Fig. S12.**
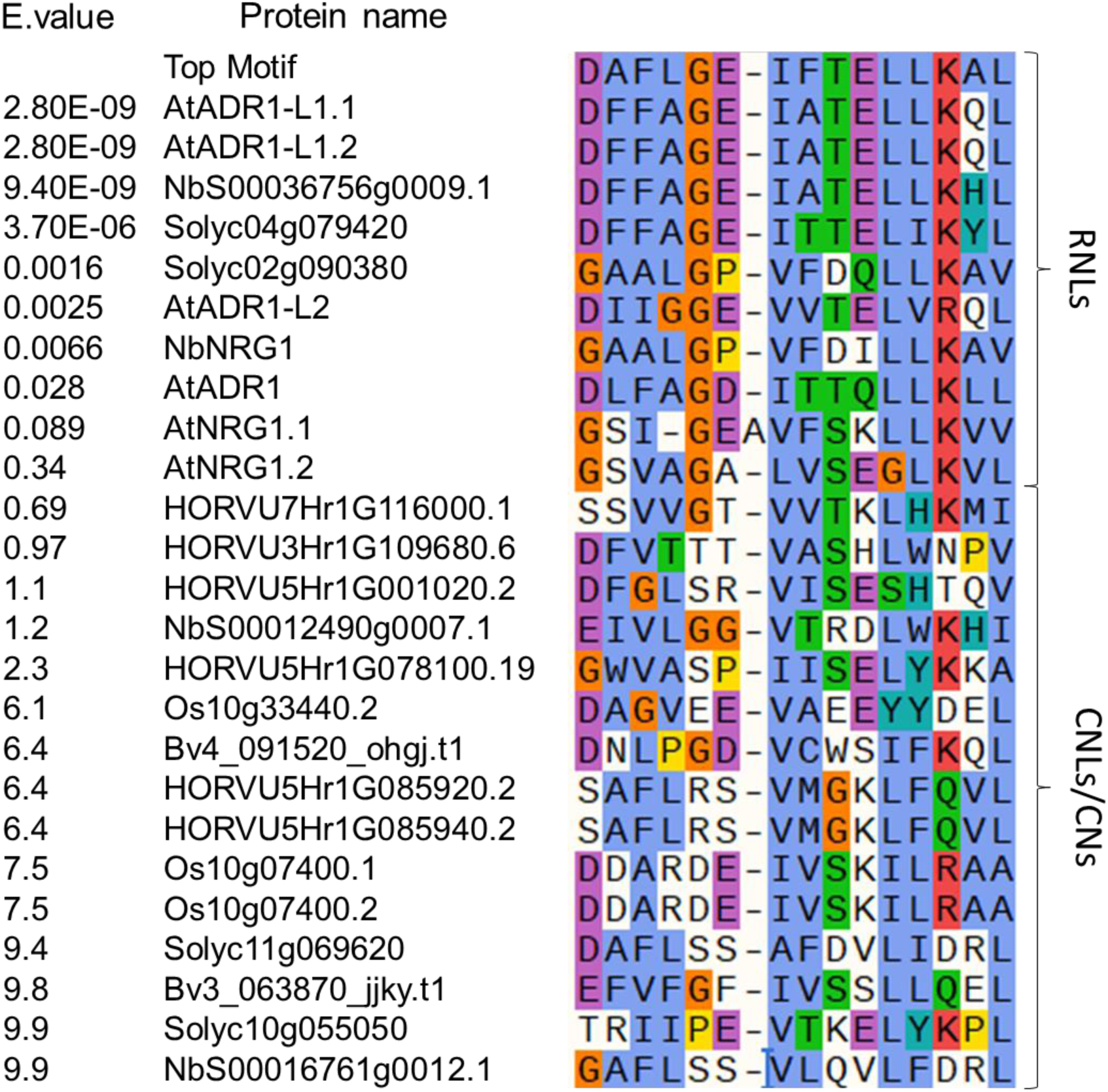
The RNL N-terminal motif is not shared by CNLs. The RNL N-terminal motif was used to scan (MAST, meme-suite.org, (*54*)) a database comprising 988 putative CC-NLRs and CC_R_-NLRs from six representative plant species (Arabidopsis, sugar beet, tomato, N. benthamiana, rice and barley) from (*55*). The top 10 motif-containing protein were all RNLs and CNLs exhibited only a very degenerated version of the motif.

**Fig. S13.**
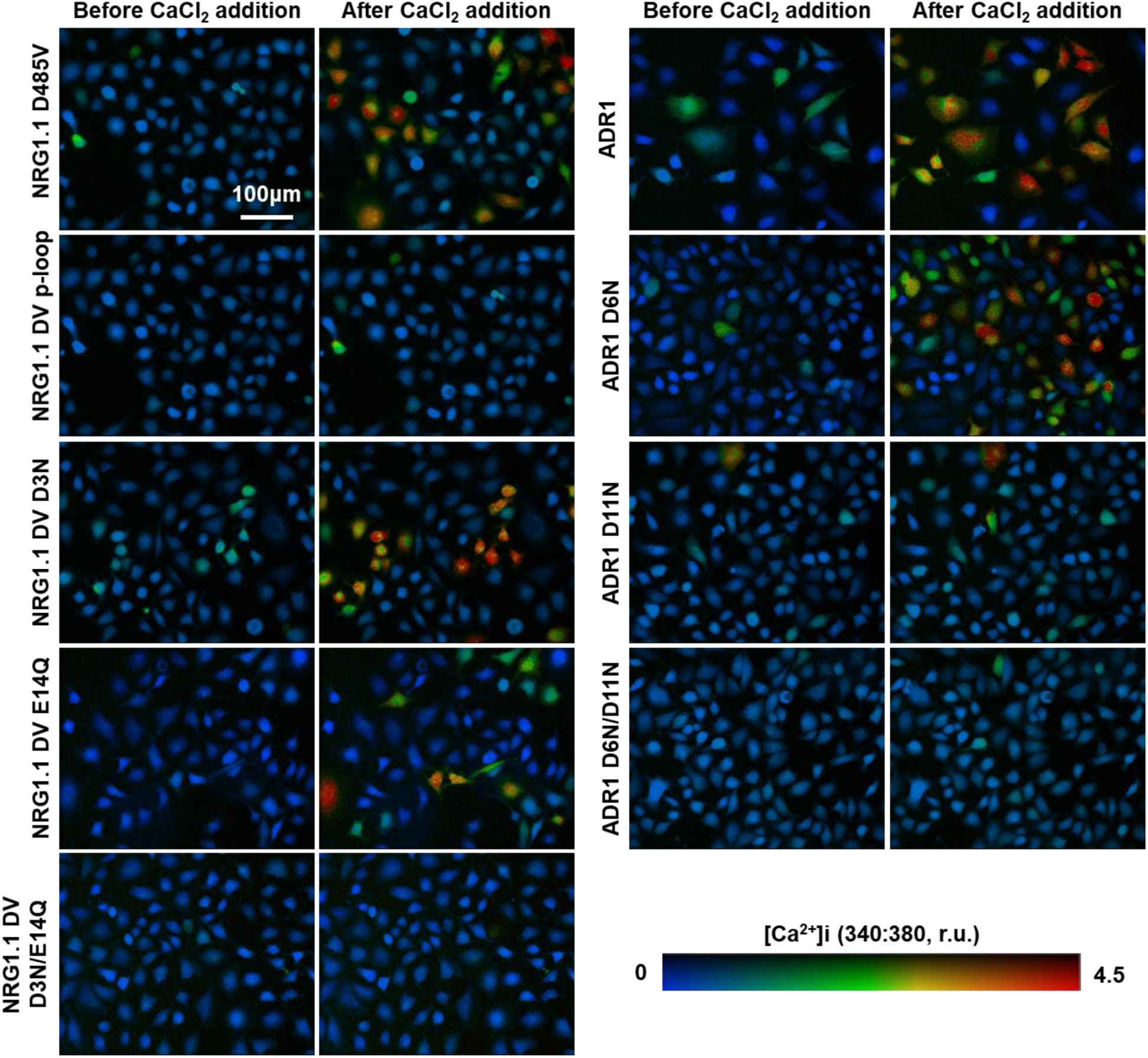
Impact of mutation in the N-terminal RNL motif on intracellular [Ca^2+^] in NRG1.1 D485V and ADR1-expressing HeLa cells, as visualized with Fura-2. Representative fluorescence microscopy images of HeLa cells expressing active NRG1.1 D484V, ADR1 or variants, before or 2 minutes after 2.5mM CaCl_2_ addition. Intracellular [Ca^2+^] was calculated from Fura-2 emission ratios (340:380 nm) and scaled using a pseudo-color bar. The ratios were quantified in Fig4 H and J.

**Fig. S14.**
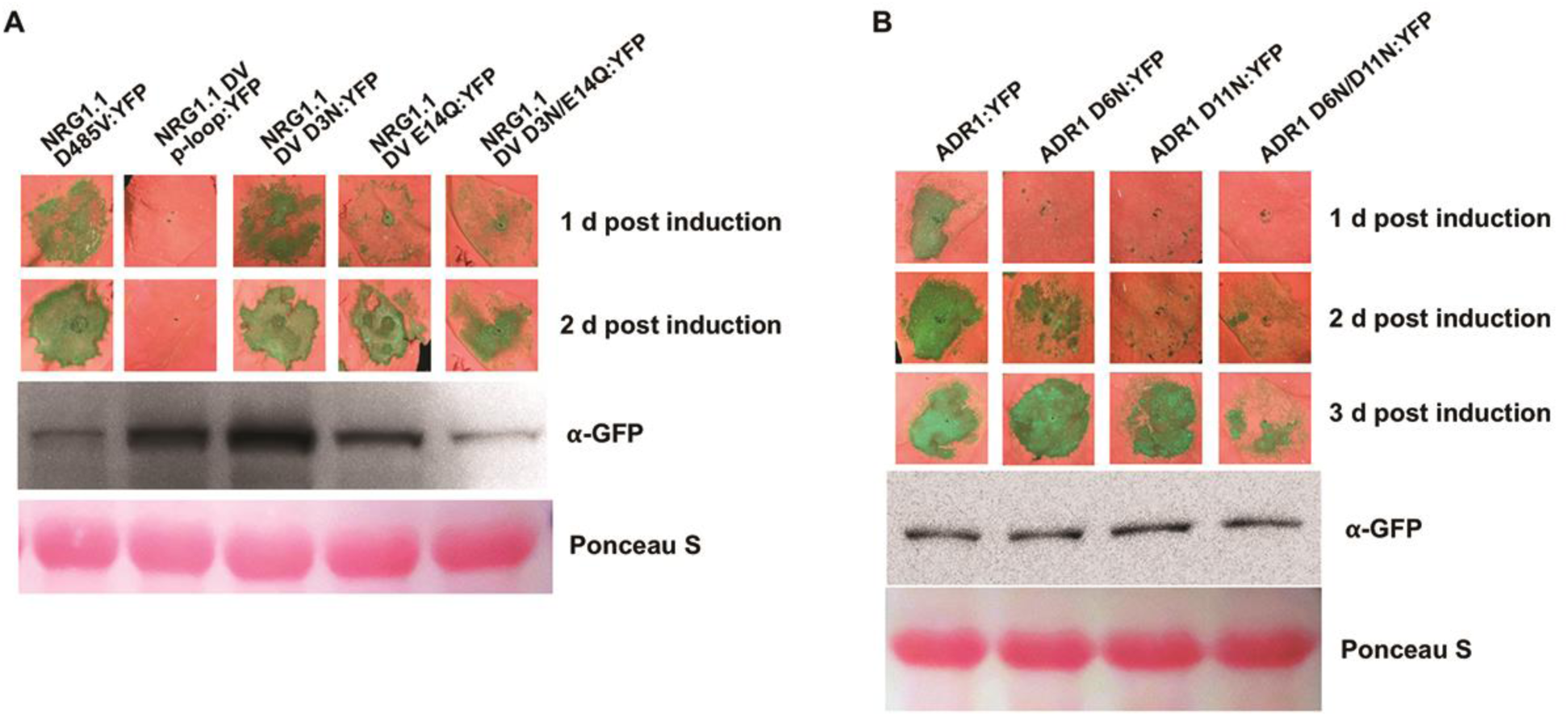
Mutation of N-terminal negatively charged residues in NRG1.1 and ADR1 delays cell death in *Nicotiana benthamiana*. UV-image of *Nb* leaves infiltrated with Agrobacterium harboring indicated constructs to observe cell death phenotypes. YFP tagged NRG1.1 and ADR1 proteins extracted from harvested tissues 6 hours post induction were resolved by SDS-PAGE and blotted with anti-GFP antibody.

**Fig. S15.**
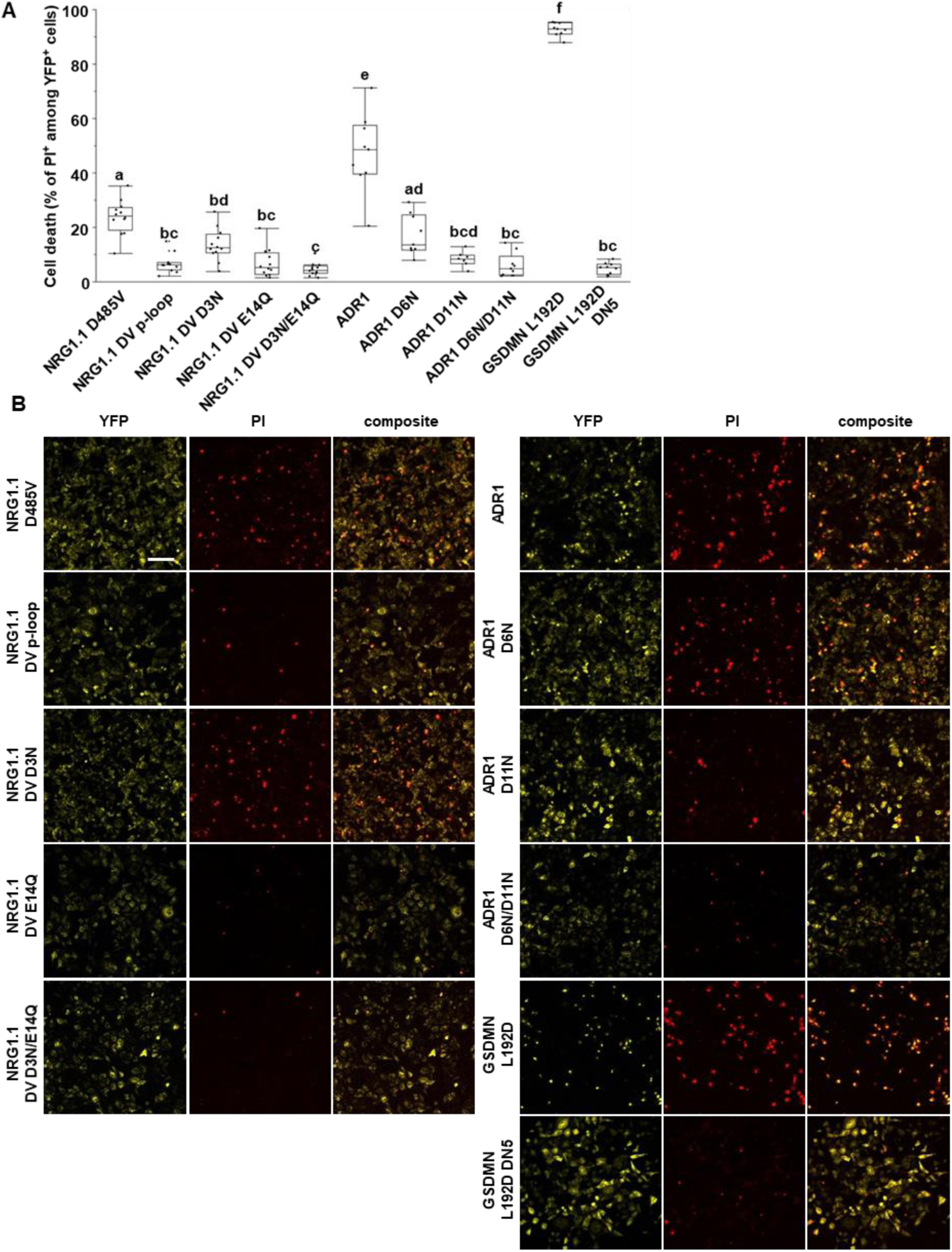
Negatively charged residues in the conserved RNL N-terminal motif are required for cell death in HeLa cells. **A** Cell death percentage observed 6 hours post-induction (hpi) of protein expression with 1μg.ml^-1^ doxycycline. Cells were stained with propidium iodide (PI) and observed with a confocal microscope. The cell death is the percentage of PI^+^ cells among the YFP^+^ cells. The data presented here comes from 3 independent experiments. Letters represent statistical difference (ANOVA with post-hoc Tukey HSD test, p-value<0.05). **B** Representative fluorescence images used for cell death quantification in A. Scale bar is 100μm.

**Fig. S16.**
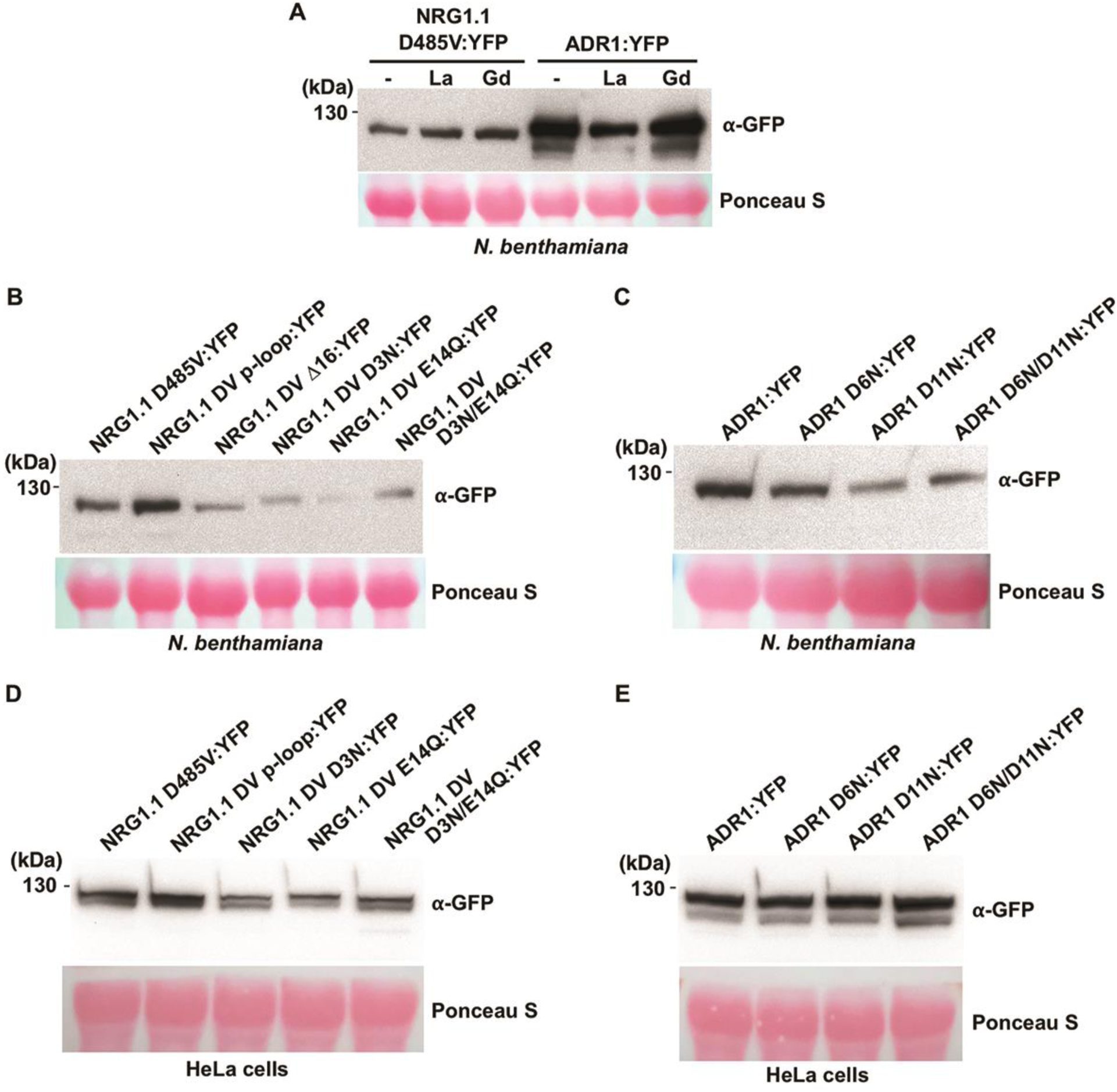
RNL proteins are correctly expressed in *N. benthamiana* and HeLa cells. Total proteins extracted from samples of Fig. 4B (**A**), Fig. 4C and 4D (**B**), Fig. 4E and 4F (**C**), Fig. 4G and 4H (**D**), and Fig. 4I and 4J (**E**) were resolved in 8% SDS-PAGE gel and transferred to nitrocellulose membranes. The RNL proteins were detected by immunoblotting the membranes with anti-GFP antibody.

**Table Sl:**
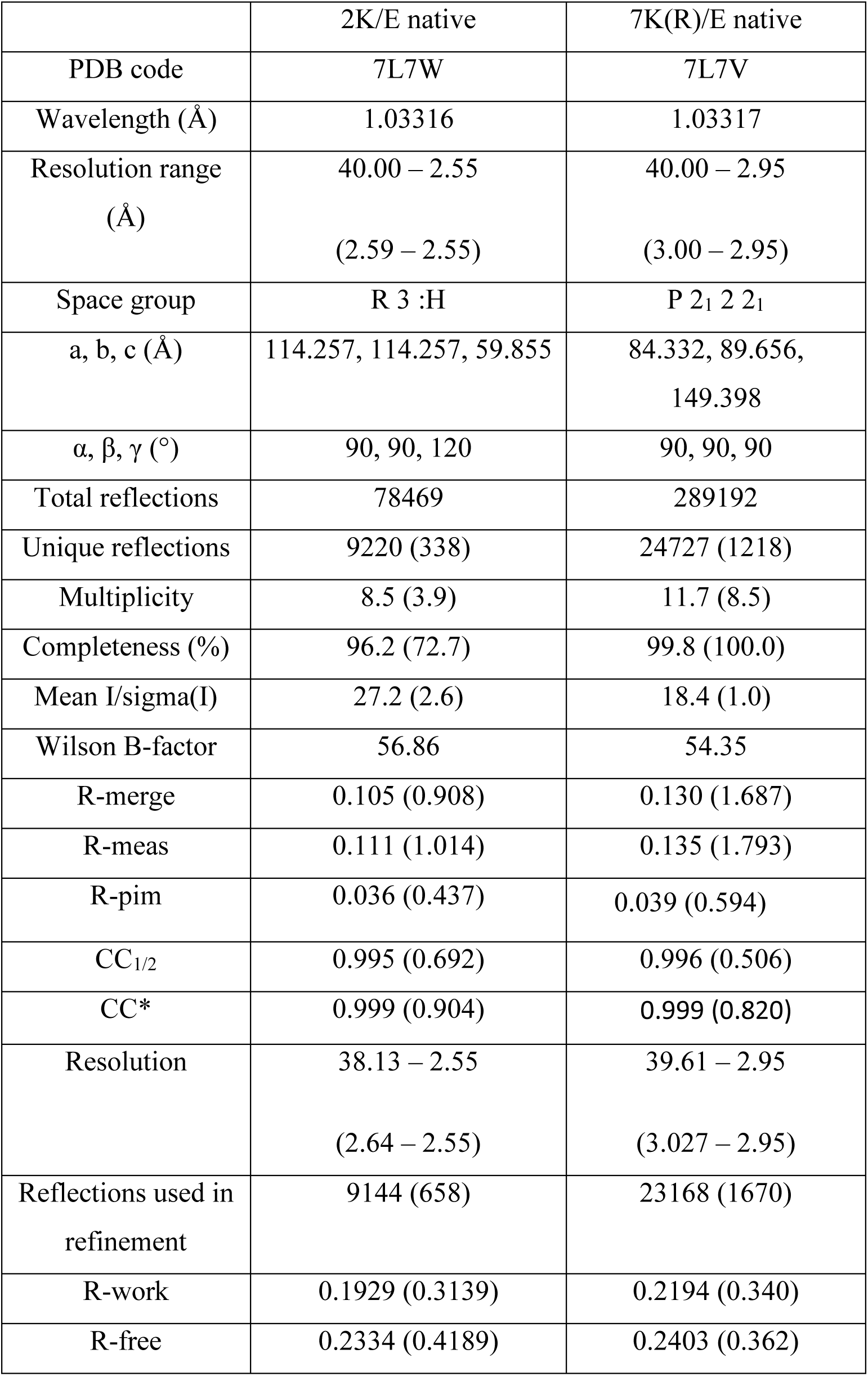
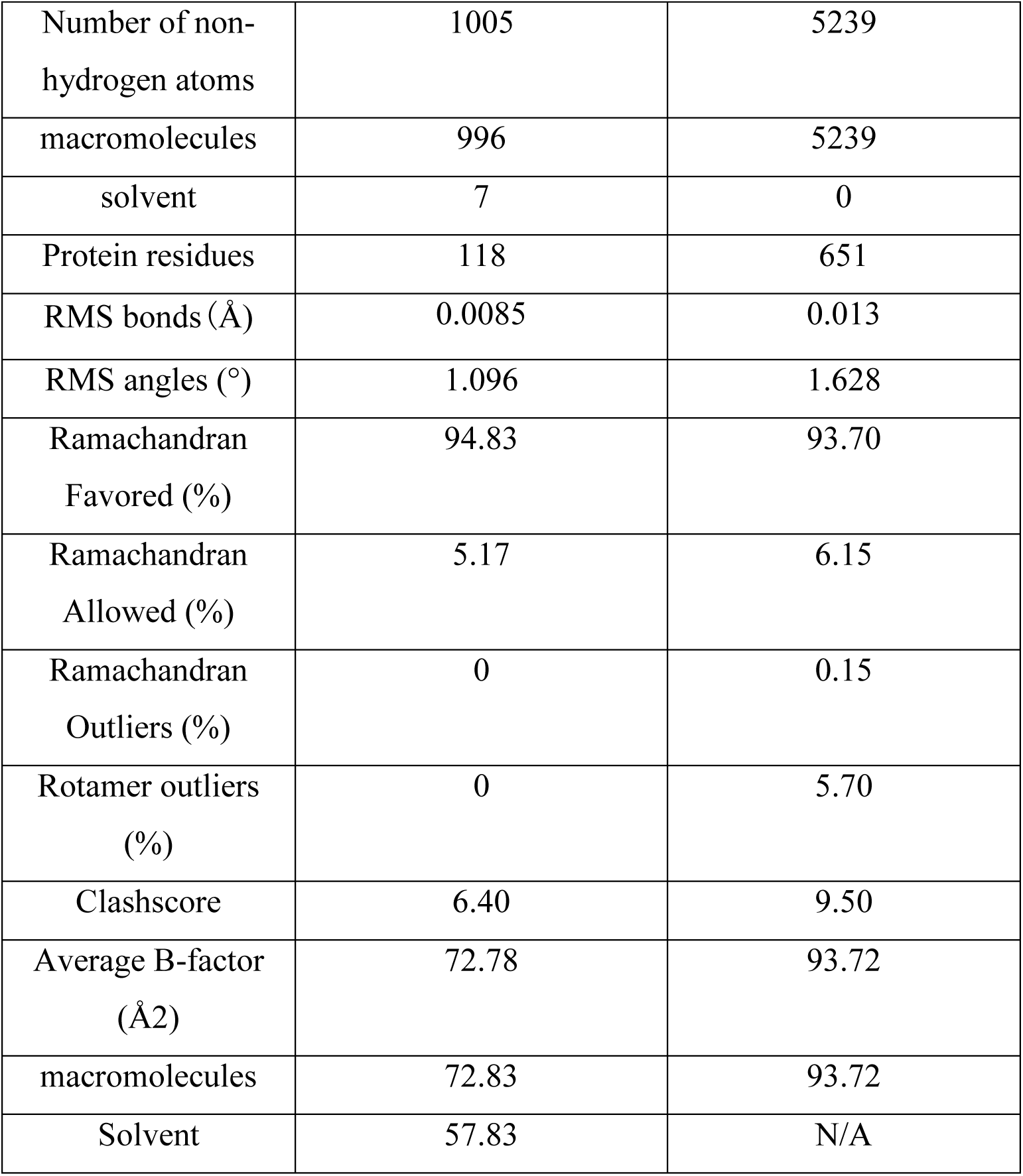
Crystallography data collection and refinement statistics

**Table S2:** Proteins structurally similar to AtNRG1.1 2K/E, based on program Dali

| PDB ID | Z-score | RMSD (Å) | Description |
| --- | --- | --- | --- |
| 6j5v | 10.0 | 2.3 | Arabidopsis CC-NB-LRR ZAR1 |
| 4m70 | 9.4 | 2.4 | Potato CC-NB-LRR Rx |
| 4btf | 9.3 | 2.4 | Mouse MLKL |
| 2ncg | 8.8 | 2.8 | Wheat CC-NB-LRR Sr33 |
| 6e7e | 8.5 | 2.7 | Inclusion membrane protein A |
| 3rip | 8.5 | 3.2 | Gamma-tubulin complex component 4 |
| 3ay5 | 8.4 | 2.8 | Cyclin-D1-binding protein 1 |
| 3zni | 8.1 | 2.7 | E3 Ubiquitin-protein Ligase CBL-B |
| 1r0d | 8.0 | 3.0 | Huntingtin interacting protein 12 |
| 5y2g | 7.9 | 3.5 | Maltose-binding Periplasmic protein, protein B |

**Table S4:**
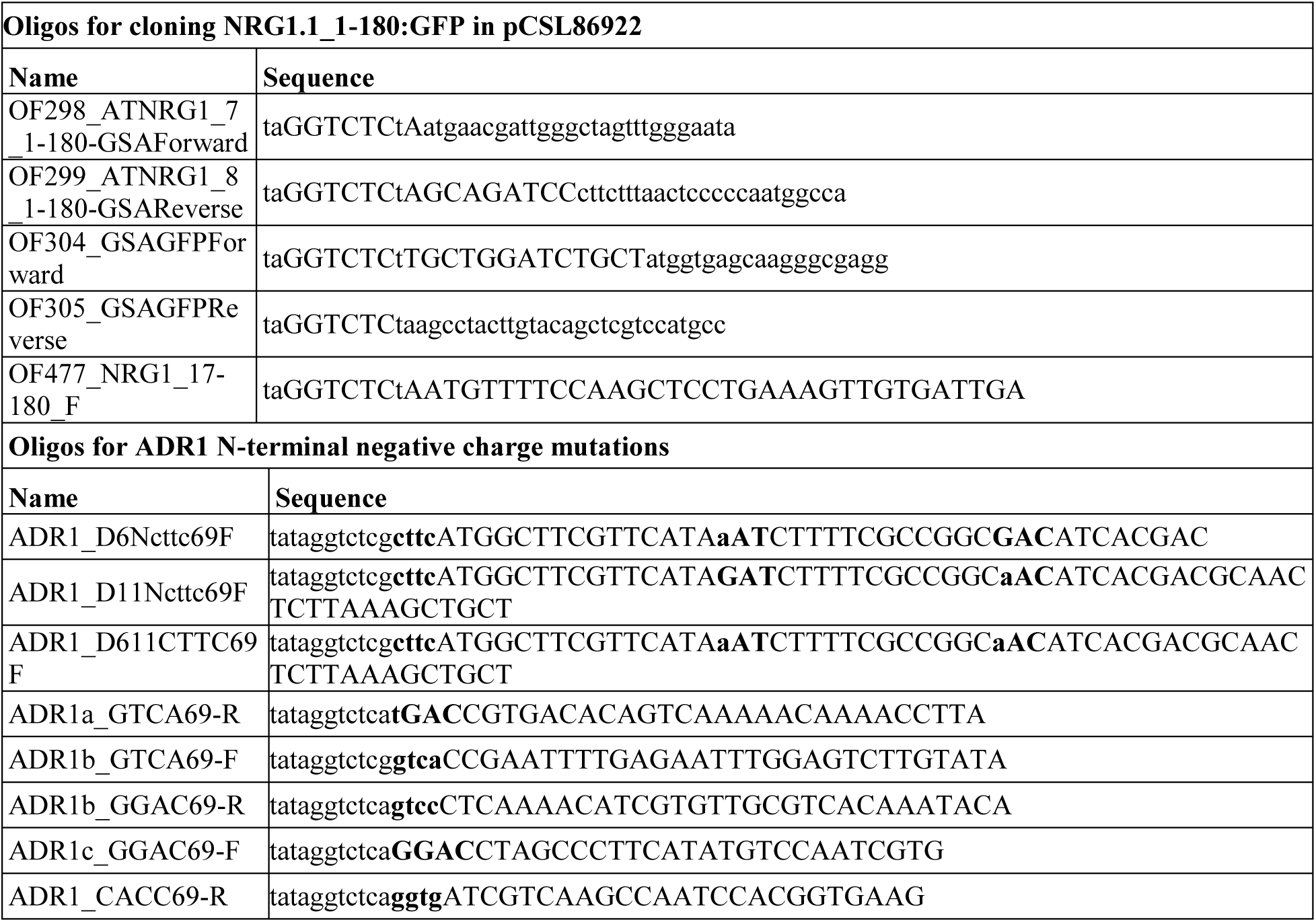
List of oligos used in this study

